# Identification of electroporation sites in the complex lipid organization of the plasma membrane

**DOI:** 10.1101/2021.10.16.464625

**Authors:** Lea Rems, Xinru Tang, Fangwei Zhao, Sergio Pérez-Conesa, Ilaria Testa, Lucie Delemotte

**Author notes:** Authors contributed equally (Xinru Tang, Fangwei Zhao).

## Abstract

The plasma membrane of a biological cell is a complex assembly of lipids and membrane proteins, which tightly regulate transmembrane transport. When a cell is exposed to a strong electric field, the membrane integrity becomes transiently disrupted by formation of transmembrane pores. This phenomenon, termed electroporation, is already utilized in many rapidly developing applications in medicine including gene therapy, cancer treatment, and treatment of cardiac arrythmias. However, the molecular mechanisms of electroporation are not yet sufficiently well understood; in particular, it is unclear where exactly pores form in the complex organization of the plasma membrane. In this study we combine coarse-grained molecular dynamics simulations, machine learning methods, and Bayesian survival analysis to identify how formation of pores depends on the local lipid organization. We show that pores do not form homogeneously across the membrane, but colocalize with domains that have specific features, the most important being high density of polyunsaturated lipids. We further show that knowing the lipid organization is sufficient to reliably predict poration sites with machine learning. However, by analysing poration kinetics with Bayesian survival analysis we then show that poration does not depend solely on local lipid arrangement, but also on membrane mechanical properties and the polarity of the electric field. Finally, we discuss how the combination of atomistic and coarse-grained molecular dynamics simulations, machine learning methods, and Bayesian survival analysis can guide the design of future experiments and help us to develop an accurate description of plasma membrane electroporation on the whole-cell level. Achieving this will allow us to shift the optimization of electroporation applications from blind trial-and-error approaches to mechanistic-driven design.

## 1 Introduction

The plasma membrane of a cell is a complex assembly of hundreds of different types of lipids and membrane proteins, which tightly regulate transmembrane trafficking and participate in cell signalling (1, 2). The molecular organization of the plasma membrane and its integrity are essential for the life of the cell. However, when the cell is exposed to external forces, the membrane integrity can become transiently disrupted by formation of transmembrane pores. Such disruption can be useful in many clinical applications, for example when nucleic acids need to be delivered across the plasma membrane into the cell interior, where they can carry out their tasks (3, 4). Various physical methods can induce transmembrane pores, including ultrasound, light, electric field, and mechanical deformation (stretching/squeezing) (5–8). In terms of clinical applications, poration by the application of electric fields or *electroporation* is the most widely used. It is approved for treatment of solid tumours, and it is being tested in clinical trials for gene therapy, vaccination against cancer and infectious diseases, and for cardiac ablation (9–12).

Despite being used in many applications in medicine and biotechnology, the molecular mechanisms underlying electroporation are not sufficiently well understood. The most accepted models, that describe electroporation on the whole-cell level, consider that pores can form only in the lipid domains of the plasma membrane and that all pores exhibit a similar kinetic behaviour (13–15). However, accumulating evidence from experiments and simulations on model systems speaks against these assumptions. Poration kinetics in pure lipid bilayers has been shown to depend on the type of lipids and their phase state (16, 17). Since the lipids in the plasma membrane organize in domains (18, 19), there must exist locations which are more and less prone to poration. Moreover, our research suggests that pores can nucleate within some membrane proteins, causing protein denaturation and lipid rearrangement (20). Such lipid/protein pores can be more stable than pure lipid pores and are more likely to explain the persistent increase in plasma membrane permeability following exposure to electric pulses. Studies have further shown that pore formation and/or expansion is affected by the actin cytoskeleton, either via actin’s influence on lipid organization or the mechanical properties of the membrane (21, 22). The current challenge is to gather this ensemble of findings into a coherent and predictive mathematical model describing electroporation of the living cell’s plasma membrane. In a living cell’s plasma membrane, pores cannot form anywhere: as soon as a sufficient number of pores is formed, the transmembrane voltage drops, preventing formation of new pores (23, 24). In other words, pores will form preferentially in specific sites with the highest poration propensity. However, it remains to be elucidated which are the properties of these sites.

The challenge of studying pores in the plasma membrane experimentally is that pores are nanometer-sized and open only transiently, whereby most of them appear to rapidly close (ns-μs range) after turning off the electric field (25–27). Pores have been imaged in real time in droplet interface bilayers with TIRF microscopy (17); however, the membranes were exposed to seconds-long electric pulses, which are much longer than pulses used in electroporation applications (ns–ms range), and which would likely not be tolerated by living cells. A few attempts have been made to visualize pores in cells using electron microscopy (28, 29); however, the observed pores were suggested to be artifacts of sample preparation (30). Overall, the current state of experimental methods does not appear to be at a stage where it would provide the spatiotemporal resolution required to understand the molecular mechanisms of plasma membrane electroporation in its entirety.

In this study we thus resort to molecular modelling methods to investigate plasma membrane electroporation. In particular, we use coarse-grained molecular dynamics simulations, building on their success in studying membrane lateral organization and dynamic behaviour (31–33). By running electroporation simulations on lipid membranes mimicking the realistic composition of plasma membranes, we confirm that pores do not form homogeneously across the membrane, but colocalize with domains that have specific features, particularly high content of polyunsaturated lipids. By training machine learning algorithms, we further demonstrate that knowing the local lipid distribution is sufficient to predict with ~80% accuracy the locations, which are most likely to be porated. However, by analysing poration kinetics with Bayesian survival analysis we then show that poration does not depend solely on local lipid arrangement, but also on membrane mechanical properties and the polarity of the electric field. Finally, we discuss how atomistic and coarse-grained molecular dynamics simulations, machine learning methods, and Bayesian survival analysis combined can help us develop more accurate cell-level models, which are required to foster new and better electroporation-based applications.

## 2 Results

To study plasma membrane electroporation we have used coarse-grained membranes consisting of >60 different lipid types parametrized with the Martini force-field (34). The membranes mimic the composition of either an idealized average mammalian plasma membrane (APM) or a human brain plasma membrane (BPM) and have been developed and equilibrated in earlier work (35, 36). The lipid composition of both APM and BPM is asymmetric, the lipids in the outer leaflet being different from those of the inner leaflet. Both compositions contain similar lipid types but differ in their fractions (Fig. 1). Compared with BPM, APM has lower fraction of cholesterol, higher fraction of phosphatidylcholine (PC) lipids and higher fraction of monounsaturated (MU) and polyunsaturated (PU) lipids in the top leaflet.

**Fig. 1:**
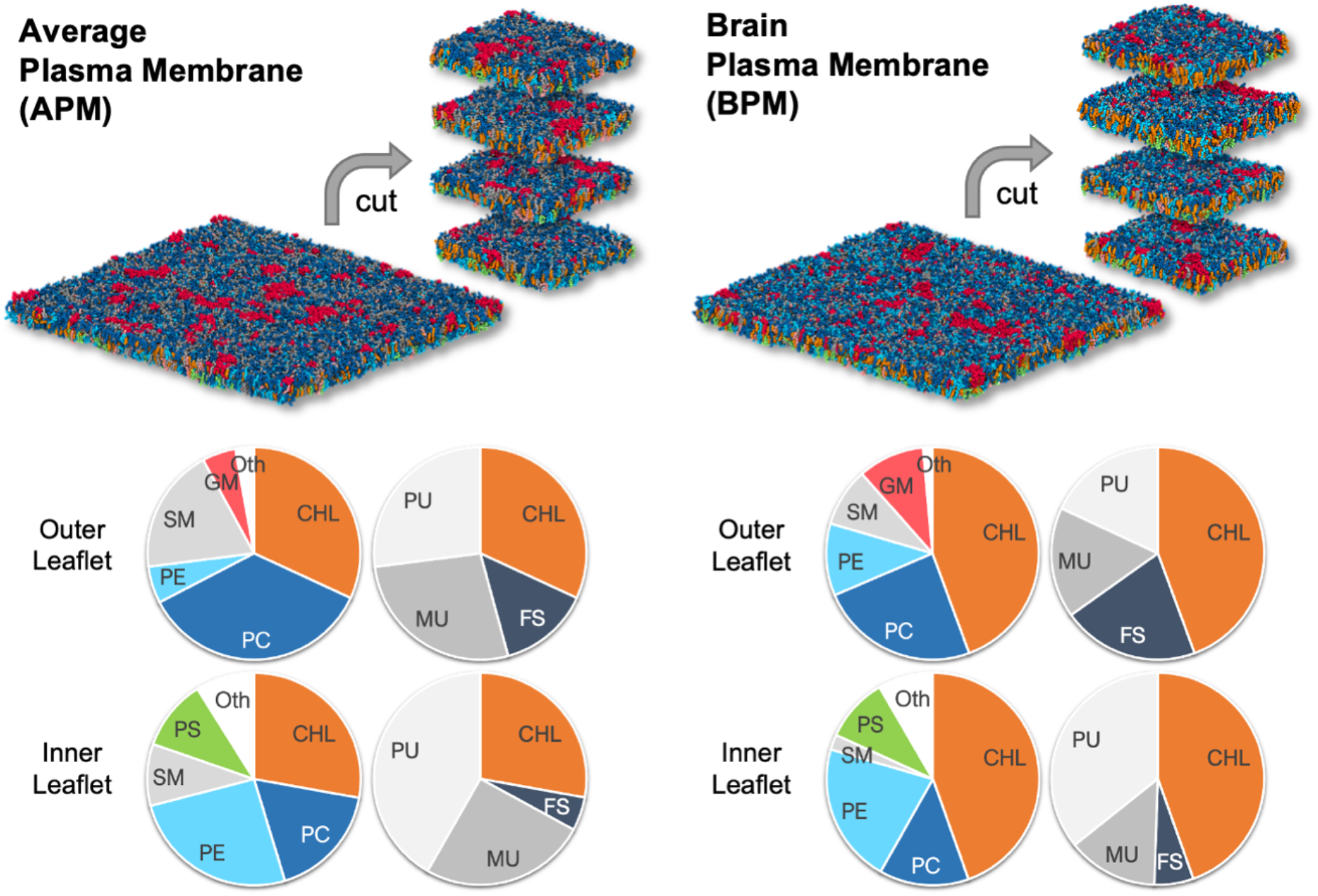
The average mammalian and human brain plasma membranes. Two equilibrated membranes taken from the study of Ingolfsson et al. (36) were cut into four 30 nm × 30 nm large pieces each. The pie charts show the fraction of lipid subgroups in the inner and outer leaflet of the membranes. Further details about the lipid composition are given in Suppl. Tables S1 and S2 and Suppl. Fig. S1. CHL-cholesterol, PC-phosphatidylcholine, PE-phosphatidylethanolamine, SM-sphingomyelin, GM-gangliosides, PS-phosphatidylserine, FS-fully saturated lipids, MU-monounsaturated lipids, PU-polyunsaturated lipids. Figure inspired by (36).

The following subsections present the results and analysis as follows. First, we present the results from electroporation simulations and demonstrate that all membranes exhibit preferential poration sites. Then we determine local membrane properties and ask which of them increases/decreases the poration propensity. We further investigate the importance of local membrane properties by training machine learning algorithms and predicting the sites that are most likely to be porated. Finally, we apply Bayesian survival analysis to investigate how membrane properties influence the poration kinetics and to develop an underlying kinetic model.

### 2.1 Membranes exhibit preferential poration sites

The original APM and BPM membranes from Ingolfsson et al. (35, 36) were cut into 4 pieces each (Fig. 1) to increase sampling, since the analysis was focused on the first poration event. After a short re-equilibration, each of the 8 membranes was subject to electroporation simulations, 60 simulations under hyperpolarizing field and 60 simulations under depolarizing electric field. Both polarities of the electric field were considered, since during exposure of a cell to an electric field, the plasma membrane becomes hyperpolarized on the side facing the positive electrode and depolarized on the side facing the negative electrode. The electric field was set to ±127.7 mV/nm, which was high enough to induce a pore within ~15 ns. Being able to observe poration over short time scales was important to minimize lateral lipid diffusion and provide a reliable mapping of local membrane features before electric field application to the likelihood of a poration event. After identifying when and where the pores formed in each simulation, we observed that pore locations do not distribute homogeneously along the membrane surface but often cluster together (Fig. 2). An individual membrane exhibits one or more of such clusters. The location of pores is similar albeit not completely identical under depolarizing and hyperpolarizing electric field.

**Fig. 2:**
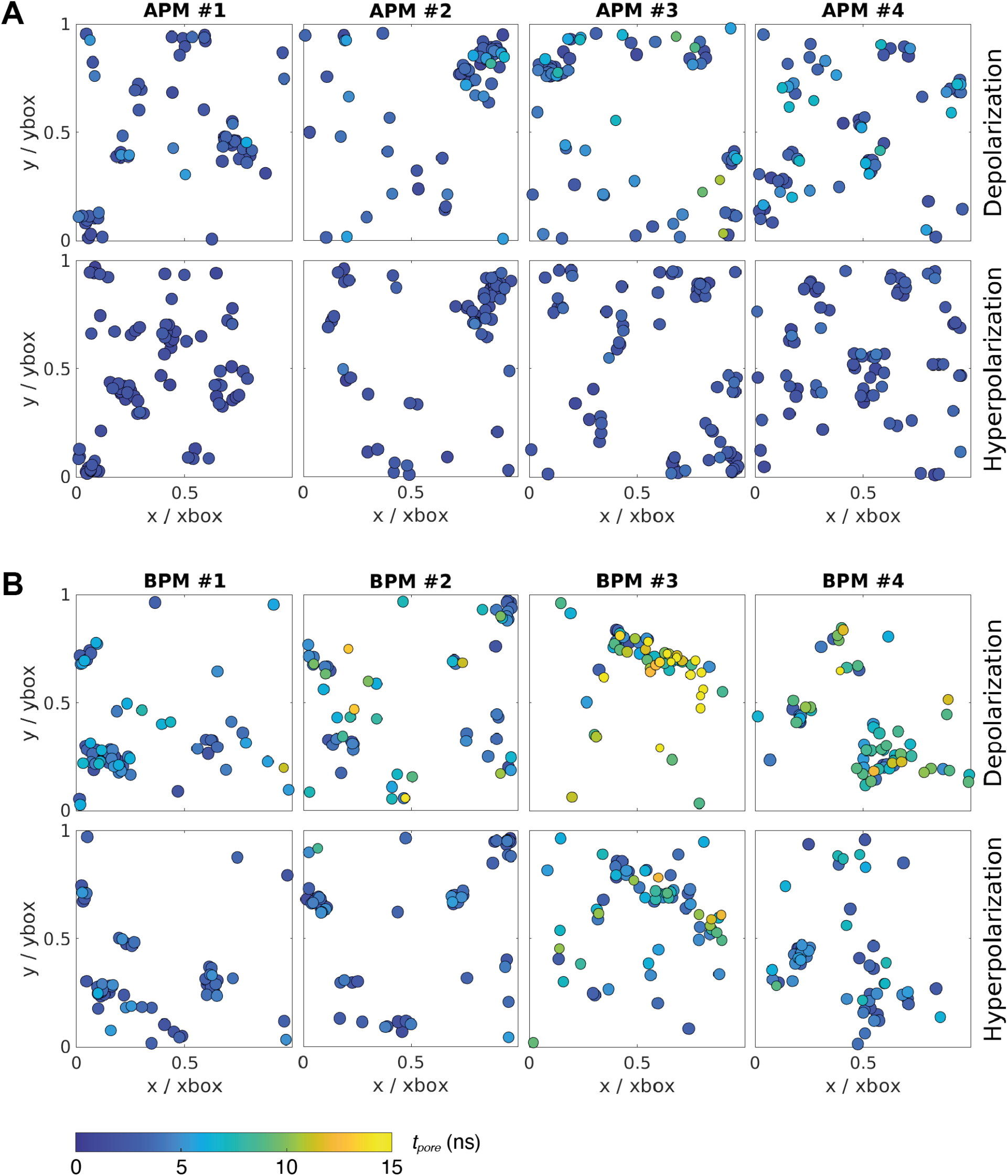
Location and kinetics of pore formation in each membrane upon depolarization and hyperpolarization. The pore locations are expressed relative to the dimensions of the simulation box at the poration time, to correct for the natural fluctuations in the membrane area during the simulation. The colour of the circle codes for poration time according to the colour bar.

### 2.2 Poration sites colocalize with domains with specific features, in particular with a high density of polyunsaturated lipids

We hypothesized that pores preferably form in nanodomains with specific features. We used the recently developed tool MemSurfer (37) to extract from each membrane the local area per lipid (APL), membrane thickness, mean curvature, cosine of the dipole angle (cos *θ*_*dip*_), charge, and lipid tail order parameter. We also determined the local density of individual groups of lipids, grouping the lipids either according to their head architecture or their tail saturation. These membrane features were extracted from 101 frames of a 10-ns long trajectory before electric field application, from each leaflet separately and in points corresponding to locations where a pore has or has not formed in any of the electroporation simulations. The first and second group of points are labelled as “porated” and “non-porated” locations, respectively (Fig. 3A, left). We estimated the distribution of the values of the various features in non-porated and porated locations by constructing histograms (Suppl. Section S5) and quantified the difference between probability density estimates by means of the Kullback–Leibler divergence (Fig. 3B). By far the most significant feature to distinguish the locations where pore formed from locations where they did not was the presence of polyunsaturated lipids, in both the APM and BPM membranes. This finding is corroborated by visualizing the colocalization of pore clusters with nanodomains enriched with polyunsaturated lipids (Fig. 3A). The analysis further showed that pores in all membranes are favoured in regions which have greater area per lipid, lower lipid order, higher density of phosphatidylcholine (PC) lipids, and lower densities of cholesterol (CHOL), gangliosides (GM) and fully saturated (FS) lipids. In BPM, the pores are also favoured in regions with higher content of phosphatidylethanolamine (PE) lipids. None of the analysed features appeared markedly dependent on the electric field polarity, even when we contrasted them with 10-ns-long trajectories obtained under non-porating electric field (Suppl. Fig. S9).

**Fig 3:**
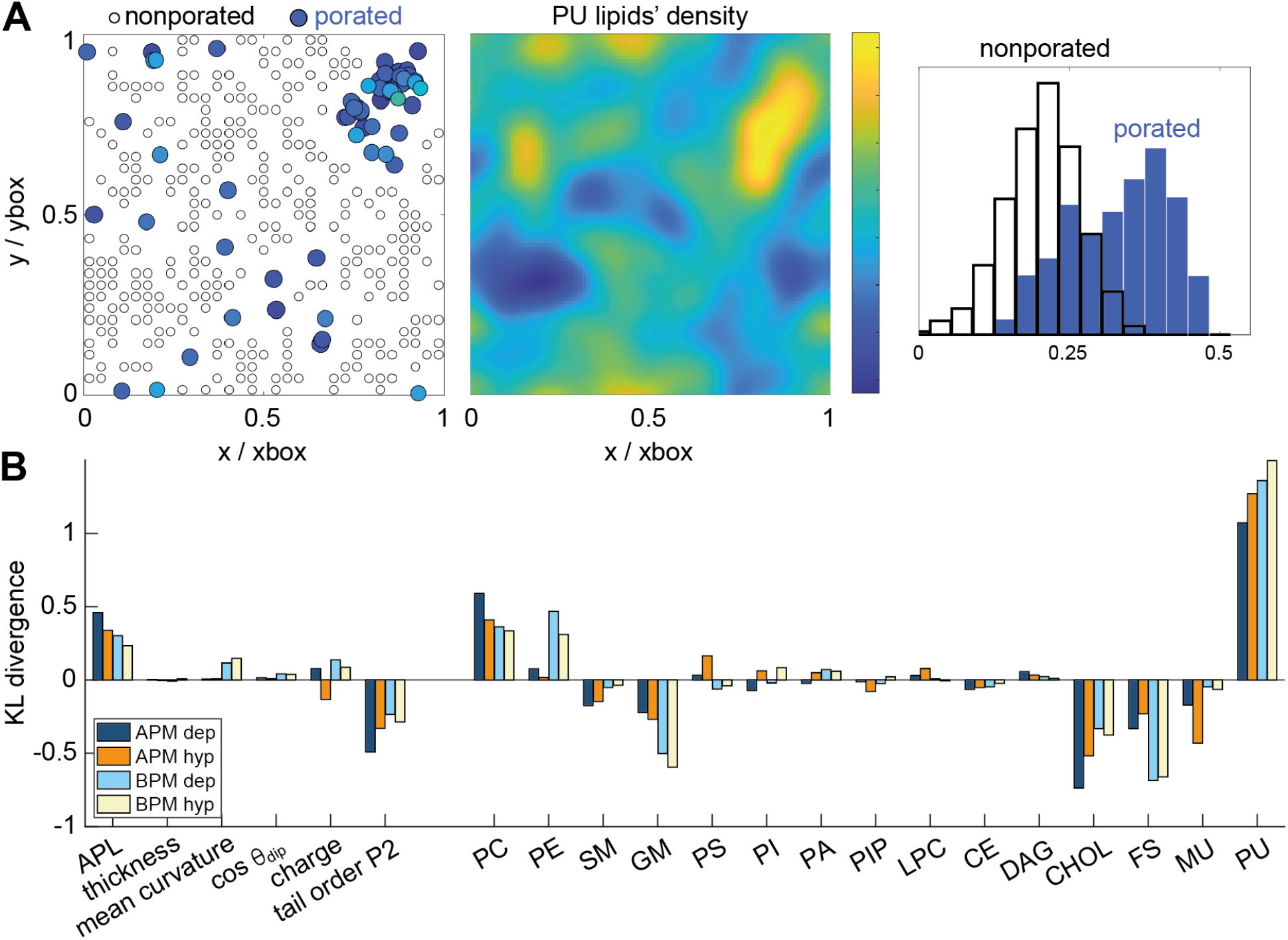
Membrane features that favour and disfavour poration. A) (*Left*) Definition of porated and non-porated locations in one of the membranes; (*middle*) most pore locations colocalize with increased polyunsaturated lipids density; (*right*) the corresponding histogram of values in porated and non-porated locations. B) Distances (KL divergence) between probability density estimates of individual features in porated and non-porated locations. The higher the bar, the more the feature influences poration. Positive and negative bars shows whether a feature favours or disfavours poration, respectively. APL-area per lipid, PC-phosphatidylcholine, PE-phosphatidylethanolamine, SM-sphingomyelin, GM-gangliosides, PS-phosphatidylserine, PI-phosphatidylinositol, PA-phosphatic acid, PIP-phosphatidylinositol phosphate, LPC-lysolipids, CE-ceramides, DAG-diglycerol, CHL-cholesterol, FS-fully saturated lipids, MU-monounsaturated lipids, PU-polyunsaturated lipids.

### 2.3 Knowing the lipid distribution is sufficient for machine learning models to reliably predict poration sites

The distances between probability density estimates suggested that the poration propensity is governed by similar features, regardless of the type of membrane and electric field polarity. To corroborate this finding further, we trained three machine learning models, namely random forest, support vector machine (SVM), and multilayer perceptron neural network (38), on selected subsets of data, *i*.*e*., using features from APM or BPM and/or features obtained under hyperpolarization or depolarization. The accuracy of predicting poration in another subset of data typically surpassed 80% in all three models tested, with random forest exhibiting slightly superior performance (Table 1). Similar accuracy was obtained regardless of whether we trained and tested the models on (i) datasets from different membrane pairs of the same composition and the same electric field polarity (columns 1-6 in Table 1); (ii) membranes of the same composition but different electric field polarity (columns 7-8 in Table 1); or (iii) membranes of different composition (columns 9-11 in Table 1). Visual inspection of the prediction showed that locations corresponding to pore clusters are reliably predicted, whereas those that are scattered away from the clusters tend not to be predicted as accurately (Fig. 4A).

**Table 1:**
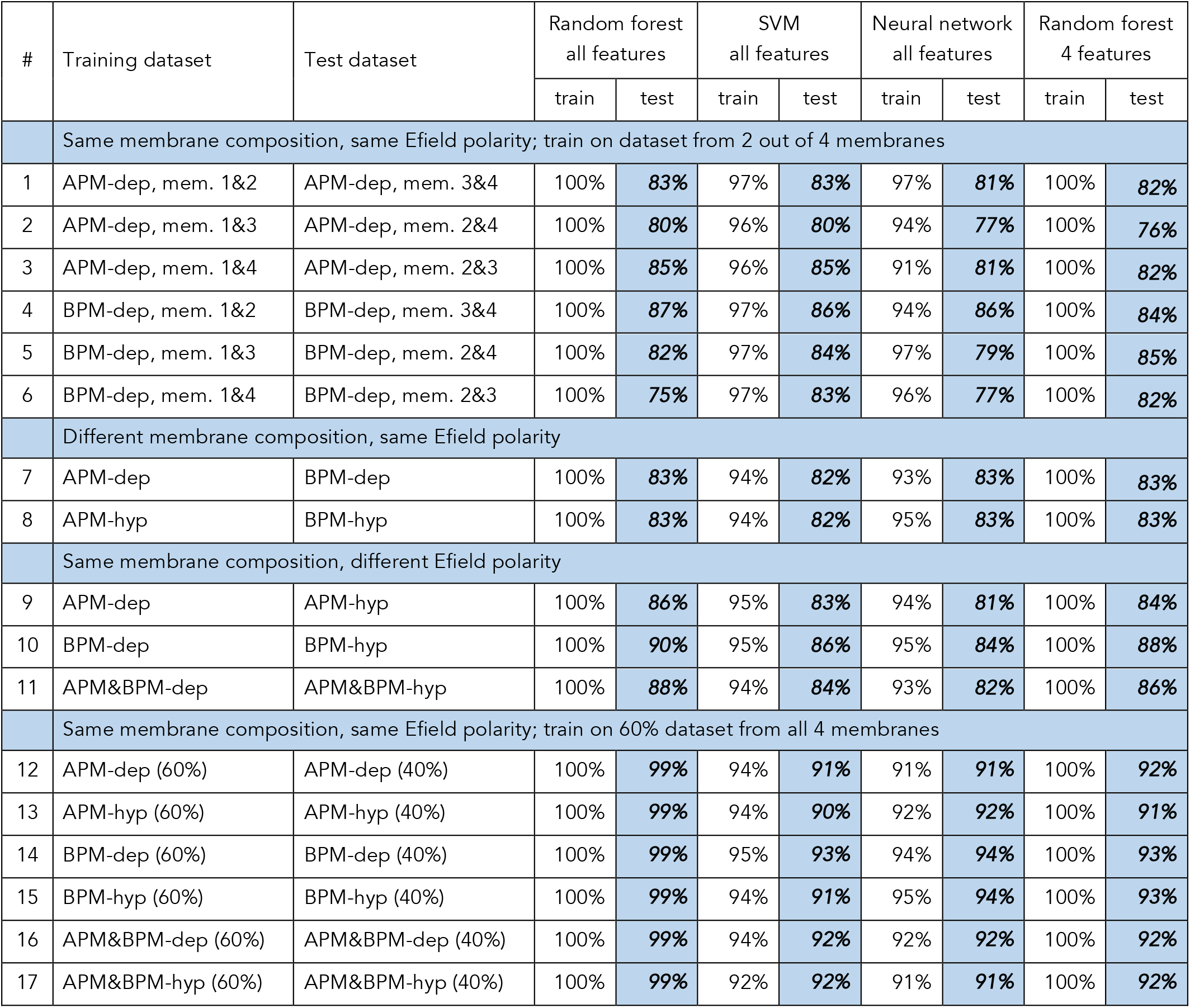
Prediction accuracy by machine learning models, reported for the training and test datasets; dep=depolarization, hyp=hyperpolarization.

**Fig. 4:**
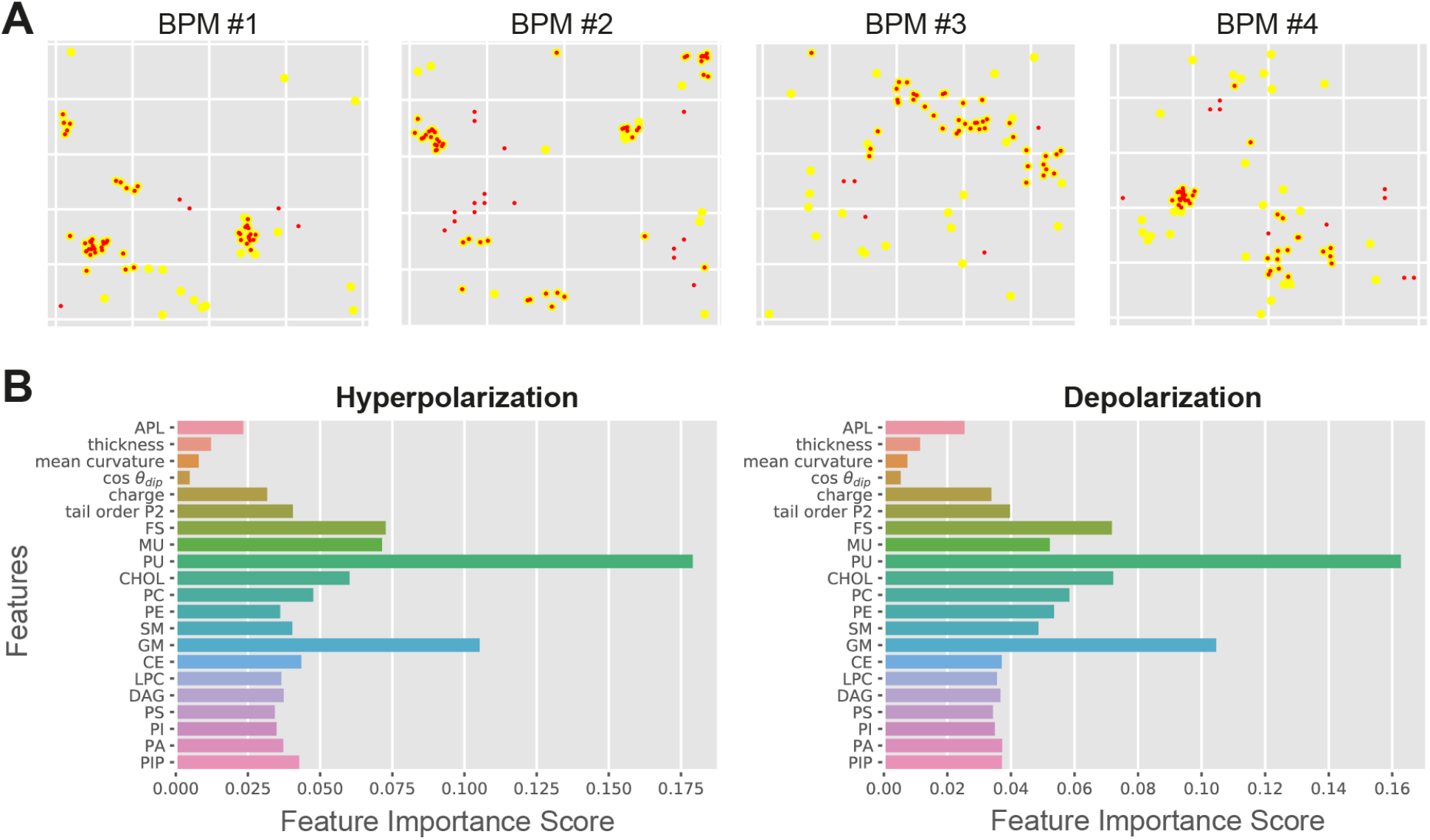
Random Forest output. A) Comparison between the predicted poration sites (red dots) and real poration sites (yellow dots). Only locations that were classified as porated in >90% of the trajectory frames are shown. B) Feature importance score for APM and BPM obtained under hyperpolarization and depolarization.

The prediction accuracy became considerably improved when a given dataset was randomly split into 60% training / 40% test dataset (columns 12–17 in Table 1). In such case, random forest reached the highest accuracy of 99%. We hypothesized that this high accuracy could be biased by oversampling, which was performed to balance the starting number of porated locations (~60) and the number of non-porated locations (300), see section 5.4. When randomly choosing 60% of a given dataset for training, the values of features from all (not just 60%) actual porated locations are effectively taken into account. Therefore, we made additional tests, where we trained random forest on the first 32 porated and 150 non-porated locations for each of the APM membrane under depolarization. The accuracy of the prediction for the rest of the porated and non-porated locations was 92%, 97%, 94%, 95% for membranes 1–4, respectively. The accuracy was lower than 99%, but still considerably higher than when training/testing on pairs of different APM membranes (columns 1–3 in Table 1). This exercise suggests that training models on data obtained from more membranes with different composition could be beneficial for predicting poration.

From the datasets for both APM and BPM, we determined the feature importance score, which confirmed that the density of polyunsaturated lipids (PU) is the most important for poration, followed by density of gangliosides (GM), cholesterol (CHOL), and fully saturated lipids (FS) (Fig. 4B). Furthermore, the feature importance score suggested that knowing the distribution of these lipid groups is sufficient to predict poration sites. The densities of other lipids and other membrane properties including the area per lipid, thickness, …, and lipid tail order can be practically neglected. We confirmed this by training random forest using 4 most important features, that is the density of PU, GM, CHOL, and FS lipids. The accuracy of the prediction was indeed very similar (≥76%) as when using all features (Table 1, last column).

### 2.4 Electroporation kinetics depends on membrane composition and electric field polarity but can be described with a universal model

The analysis presented so far demonstrates that the pore location mainly depends on the local lipid arrangement, particularly the density of polyunsaturated lipids, regardless of the type of membrane and electric field polarity. However, this does not necessarily mean that poration is equally likely or equally fast in all membranes. To investigate whether the membranes differ in poration kinetics we compared the distributions of first poration times for each membrane. These distributions were indeed found to depend both on membrane composition and electric field polarity (Fig. 5A). APM membranes tend to porate faster than BPM, and hyperpolarizing fields tend to porate faster than depolarizing fields.

**Fig. 5:**
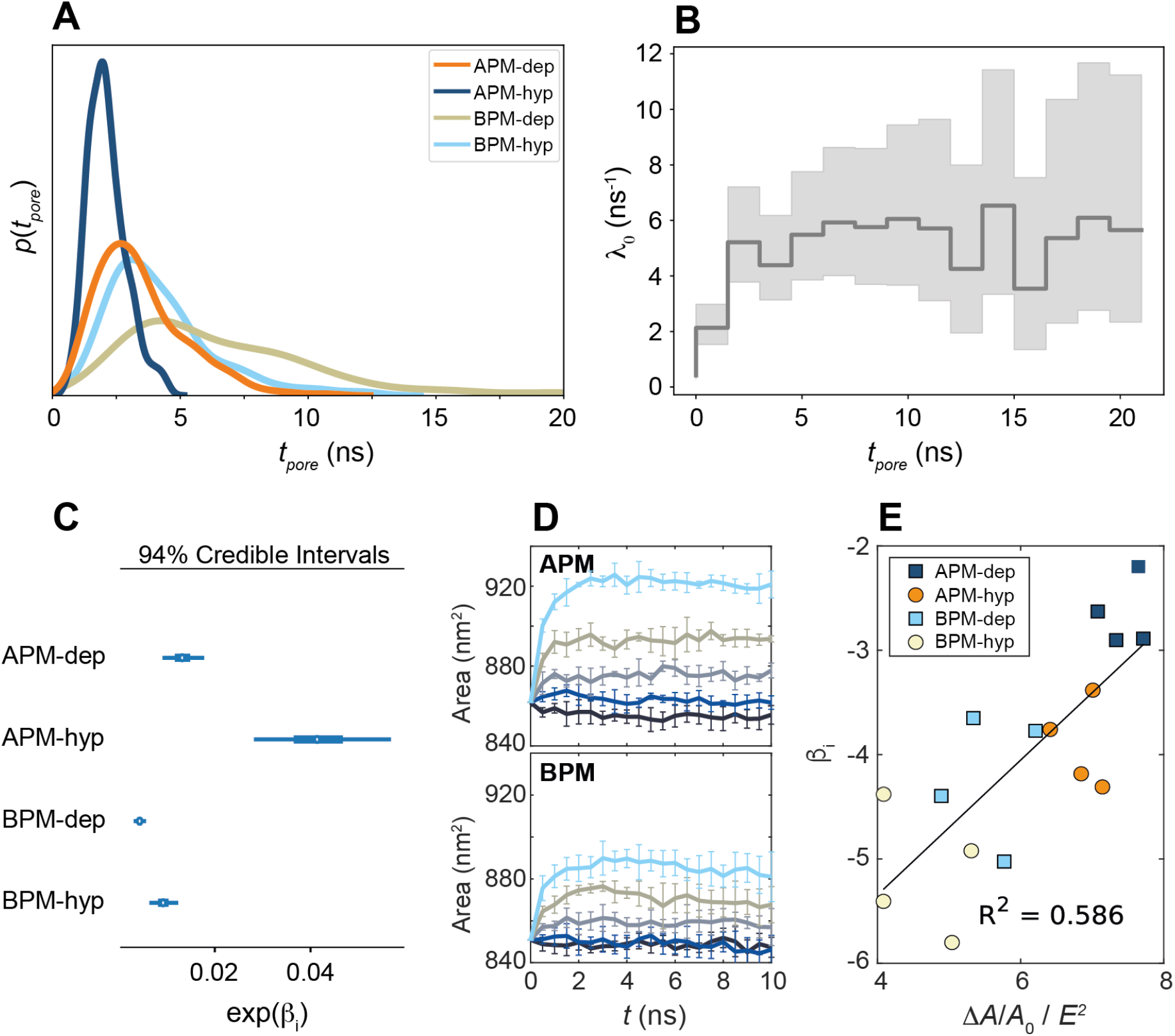
Kinetics of membrane poration. A) Probability distribution of the first poration times, sampled from 60 simulations, and approximated with kernel density estimation. B) The time course of *λ*_0_ (*t*). C) Credible intervals of exp(*β*_*i*_) for APM and BPM under depolarization and hyperpolarization. D) The change in the membrane area with time when applying non-porating electric fields *E* of 0, 42.5, 63.8, 85.1, or 106.3 mV/nm. Each curve shows mean ± s.d. of 5 simulation runs. E) Correlation between *β*_*i*_ and the steady state relative change in the membrane area Δ*A*/*A*_*0*_ normalized by *E*^2^ (Suppl. Fig. S6). The ratio Δ*A*/*A*_*0*_ / *E*^2^ is inversely proportional to the area compressibility modulus (40).

A Bayesian survival model was then trained to (i) quantify and study the statistical significance of the differences in poration rates, (ii) to investigate if an underlying universal model can be used to describe the poration kinetics, and (iii) to parametrize the kinetic parameters. Cox’s proportional hazards survival model (39) is a suitable model for this purpose, since it allows for an arbitrary time dependence of the poration rate, and it allows to then compare different systems or situations. This model assumes that for a given system, *i*, the instantaneous probability of an event happening, *λ*_*i*_(*t*) (the rate pore formation), is a function of time:

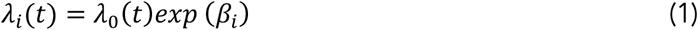

where *λ*_0_ (*t*) is a time-dependent and system-independent baseline rate of pore formation and *β*_*i*_ are regression coefficients accounting for the difference in poration kinetics between systems (APM-depolarization, APM-hyperpolarization, BPM-depolarization, BPM-hyperpolarization). The baseline rate captures the common time-dependency whereas the regression coefficient captures the system specificity. The inferred model was able to successfully reproduce the measured data (Suppl. Fig. S3) highlighting its robustness and suggesting the existence of a common kinetic model for electroporation. To further test the model validity, we trained a more complex model which allowed the *β*_*i*_ coefficients to vary with time, allowing for a system-specific time variability of (*λt*). The *β*_*i*_ obtained were found to be fairly time-independent (Suppl. Fig. S4), validating the use of the simpler model with constant *β*_*i*_.

The baseline poration rate *λ*_0_(*t*) is initially zero and then increases, reaching a constant steady state value after approximately 2.5 ns (Fig. 5B). To determine the mechanistic basis for the initial transient kinetic regime, we performed additional simulations at non-porating electric fields. These simulations revealed that the electric field *E* causes the membrane to thin and expand its area, with the relative change in membrane area proportional to *E*^2^ (Suppl. Fig. S6), as expected for Maxwell stress (41). The membrane area reaches a steady state ~2.5 ns after electric field onset (Fig. 5D), suggesting that the timescale of the initial transient kinetic regime is related to the progressive expansion of the membrane area caused by Maxwell stress.

The calculated *β*_*i*_ credible intervals (Fig. 5C) confirm that the differences in poration kinetics between different systems are statistically significant. APM membranes porate ~5 and ~2 times faster than BPM for hyperpolarizing and depolarizing fields, respectively. Hyperpolarizing fields porate membranes ~3 and ~2 times faster than depolarizing fields for APM and BPM membranes, respectively. We found that the values of *β*_*i*_ correlate with the ability of a membrane to expand its area under electric field, *i*.*e*., they corelate inversely with the membrane area compressibility modulus (Fig. 5E).

## 3 Discussion

In this study we focused on lipid pores formed by electric fields (electroporation) and we asked three main questions: Where do pores form in membranes with realistic plasma membrane lipid composition? Which membrane features/properties govern the most likely poration sites? Which membrane features/properties govern the poration kinetics?

### 3.1 Features important for membrane poration

Research conducted so far has mainly focused on studying formation of pores in model membrane systems with up to 3 different lipid types. In membranes containing a single type of fluid phase lipids, poration was found to depend on both the head and tail architecture. Longer tail length, larger headgroups (e.g., heads containing sugar moieties), and stronger intramolecular interactions between both lipid tails and heads generally reduce the propensity for poration (42– 46). Poration was further found to be more difficult in membranes with gel phase lipids compared fluid phase lipids (47–49). In binary mixtures of fluid and gel phase lipids, experiments suggested that pores form in the fluid domains (50). Mixing a lipid with cholesterol was found to either decrease or increase the poration propensity, whereby this appears to depend on the cholesterol concentration and the architecture of the lipid (51–54, 16). Studies were also done on ternary mixtures containing cholesterol, where the lipids organize in the liquid-ordered and liquid-disordered domains. Both MD simulations and experiments showed that pores preferentially form in liquid-disordered domains (55, 17).

The findings from simple model systems are in many ways reflected in our observations from membranes composed of >60 different lipid types mimicking the realistic composition of plasma membranes. We observed that pores form preferentially in locations with lower lipid tail order and larger area per lipid, and that pores avoid regions enriched with gangliosides containing large sugar headgroups, regions enriched with cholesterol and fully saturated lipids. A new finding from our study is that, in plasma membranes, the most important factor governing poration propensity is the local density of polyunsaturated lipids, regardless of the membrane type (APM or BPM) and polarity of the electric field.

### 3.2 Polyunsaturated lipids and lipid oxidation

The finding that poration is strongly facilitated in the presence of polyunsaturated lipids is very important from the standpoint of lipid oxidation. Polyunsaturated lipids are highly prone for oxidation by free radicals, whereby upon oxidation a part of the lipid tail becomes hydrophilic making the membrane considerably more permeable to ions and hydrophilic molecules (56–58). Several experimental studies have shown that electroporation is associated with oxidative lipid damage, and it is believed that lipid oxidation plays an important role in increased membrane permeability after electric field exposure (59, 60). Our study suggests that most pores form in domains enriched with polyunsaturated lipids. If pores somehow act as precursors for lipid oxidation, such as by improving the access for free radicals to lipid tails, this could provide a clue on how electroporation is related to lipid oxidation.

### 3.3 Electroporation kinetics

Bayesian survival analysis demonstrated that all membranes are characterized by two distinct kinetic regimes. Initially, no poration occurs, but progressively as the membrane area expands due to Maxwell stress, poration becomes easier and the poration rate increases, plateauing at a maximum rate when the membrane has reached its steady-state area. The steady-state poration rate was found to be inversely correlated with membrane area compressibility modulus – a measure of how resistant the membrane is to compression or expansion (Fig. 5E). Since membranes are practically volume-incompressible, the increase in area is directly related to a decrease in thickness. Membrane thinning is expected to facilitate pore formation, as water molecules need to travel shorter distance when bridging the membrane. Note that when comparing properties in porated and non-porated locations (Fig. 3B) the variation in local membrane thickness was not large enough to discriminate porated locations. However, pore formation was indeed found to be favoured in regions with larger area per lipid.

In the steady state kinetic regime, the poration rate becomes constant and the time distribution of the poration events becomes exponential. A constant poration rate and an exponential distribution of first poration times is characteristic of a Poisson behaviour of the number of pores formed per unit time. Indeed, the most accepted kinetic models of electroporation assume a constant poration rate at a fixed value of the transmembrane voltage (or the electric field within the membrane) (61, 62), which is consistent with our model in the steady-state regime. Observing the non-steady state transient kinetic regimes is beyond the time resolution of most experiments. As such, our model has the advantage of characterizing the transient initial kinetic regime, which should be important when exposing cells to increasingly-used pulses with duration in the (sub)nanosecond range (63, 64).

Electroporation rate on the whole-cell level is typically modelled as (61, 62, 65):

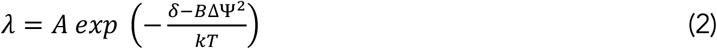

where ΔΨ, is the transmembrane voltage, *δ* is the energy barrier for pore formation at ΔΨ = 0 V, *B* is a proportionality constant, *k* is the Boltzmann constant and *T* is the temperature. *A* is a prefactor that is proportional to the number of possible pore nucleation sites and the frequency of lateral lipid fluctuations. Comparing the ratios of the pore formation rates of two systems i and j in eqs. (1-2), assuming *A*_*i*_ ≈ *A*_*j*_ and a steady-state kinetic regime we obtain:

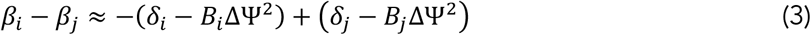

Therefore, the difference between *β*_*i*_ of two systems is approximately equal to the negative difference of their steady state poration free energy barrier in kT units. This relates the *β*_*i*_ regression coefficients to their physical interpretation. Assuming approximately equal prefactors *A*_*i*_ ≈ *A*_*j*_ is reasonable for our membranes because all membranes have the same total area. In addition, the sites with the highest poration propensity are comprised of similar lipids, so we can safely assume that those regions have similar lipid fluctuation frequency. Note that the pore formation barrier *δ* can be independently determined by free energy methods (66), whereas the parameter *B* can be inferred from simulating electroporation at different values of the transmembrane voltage; however, both these approaches are computationally more demanding. Bayesian survival analysis thus offers an alternative and simpler way to obtain parameters of eq. (2) for different systems.

### 3.4 Kinetic differences between average and brain plasma membranes

The average plasma membranes (APMs) exhibit shorter poration times compared to the brain plasma membranes (BPMs). Bayesian survival analysis confirmed that the poration kinetics in APMs and BPMs is statistically different. Compared with BPMs, APMs contain a smaller fraction of cholesterol and fully saturated lipids and a greater fraction of polyunsaturated lipids, all favouring poration and likely increasing the poration rate. In experimental studies, cells of different types, or even cells in different phases of the cell cycle, have been found to exhibit different electroporation thresholds (i.e., different electroporation propensities) (67–69). According to our results, this difference in thresholds is, at least in part, related to the fact that cells of different types can have considerable differences in their lipid composition, which further changes along the cell cycle (70, 71).

### 3.5 Kinetic differences between hyperpolarizing and depolarizing electric field

When a cell is exposed to an electric field, its membrane becomes hyperpolarized on the side facing the positive electrode (anode) and depolarized on the side facing the negative electrode (cathode). Experiments have shown that plasma membranes of different types of cells can become more permeabilized either on the anodic or cathodic side, whereby this asymmetry in permeabilization is still not completely understood (72–75). Our results suggest that the asymmetric lipid composition present in all mammalian plasma membranes favours pore formation on the anodic side, which is hyperpolarized. We found that hyperpolarization induces more profound membrane thinning compared to depolarization, consequently increasing the poration rate. Greater membrane thinning could be associated with electrophoretic drag of negatively charged lipids, which are mainly present in the inner leaflet and are pulled towards the membrane interior by hyperpolarizing electric field. In our previous study, in which we characterized pores that formed in voltage sensors of sodium voltage gated channels (20), we also observed that such pores are more easily formed under hyperpolarization, albeit this was associated with asymmetric distribution of charged protein residues. Nevertheless, there are numerous types of membrane proteins in the plasma membrane, some of which might become denaturated more easily under depolarization. Whether the cell is permeabilized more on the anodic or cathodic side might therefore depend on the plasma membrane’s lipid-protein content and the preferential sites of pore formation. This exemplifies the need to identify preferential poration sites in membranes with complex organization, which we discuss further in Section 3.6.

### 3.6 Towards building accurate cell-level models of electroporation

The cell membrane is a complex organization of lipids and proteins, whereby insights from atomistic molecular dynamics simulations suggest that the electric field can form pores both in the lipid domains and within some membrane proteins. The limitation of molecular dynamics simulations is that they are only able to model a small part of the membrane, and that they are not able to take into account the dynamic changes in the transmembrane voltage, which are present when electroporating whole cells (76, 77). Namely, on the whole cell level, the induced transmembrane voltage varies with position on the membrane and depends on the spatial profile of the membrane conductivity. As sufficient number of pores form in the membrane, they increase the membrane conductivity to the extent that starts decreasing the transmembrane voltage and prevents formation of additional pores. In other words, in molecular dynamics simulations it is possible to observe poration of practically any membrane; however, in a real cell membrane only the sites with the highest poration propensity can be porated.

To understand electroporation of living cells, we need to be able to develop equations describing the poration rate of different types of pores (pores in the lipid domain, different membrane proteins) and embed them into a system of ordinary and partial differential equations that describe electroporation on the whole-cell level (23, 24). Such models can then be used to study the increase in membrane conductivity, transmembrane molecular transport of different types of molecules, and changes in the membrane resting potential and/or action potentials induced by different parameters of electric pulses. Electroporation can be caried out with different pulse waveforms, where the duration of individual pulses can range from a few 100 picoseconds to tens of milliseconds. Exploring the pulse-parameter space *in silico* instead of through trial-and-error experimental approaches will facilitate optimization of electroporation-based applications *in vitro* and *in vivo* (78).

We envision that by combining atomistic and coarse-grained molecular dynamics simulations, machine learning methods and Bayesian survival analysis we will be able to improve existing kinetic models describing electroporation on the whole-cell level. Coarse-grained simulations, together with more mesoscopic models and experiments, are in the future anticipated to enable modelling of whole plasma membranes providing their detailed molecular organization (79). An exciting finding from our study is that knowing the lipid distribution is sufficient for identifying the most likely poration sites in lipid domains by using machine learning methods. As such, we anticipate that we will be able to use machine learning to estimate the membrane area, which is most amendable to poration, and hence estimate the prefactors in eq. (2). By performing electroporation simulations on selected membrane regions and applying Bayesian survival analysis, we can characterize the poration kinetics and simplify the determination of the corresponding kinetic parameters. For example, by following these approaches using coarse-grained membranes associated with actin filaments (80) we can investigate and quantify how the presence of actin cytoskeleton influences poration kinetics, either by affecting the local distribution of lipids or by influencing the mechanical properties of the membrane or both.

Coarse-grained simulations have their disadvantages. At present, coarse-grained simulations cannot be used to study poration of membrane proteins, as the protein secondary structure typically needs to be constrained for the protein to remain stable (81). Nevertheless, poration of membrane proteins and its corresponding kinetics can be inferred from atomistic molecular dynamics simulations (20). Furthermore, coarse-grained lipid bilayers are known to be more difficult to porate than corresponding atomistic bilayers, likely due to coarse-graining of multiple water molecules into a single particle (66). Despite being more difficult to porate, coarse-grained systems are able to represent the differences in the energy barriers for pore formation in bilayers composed of different lipid types as well as the influence of the membrane mechanical properties on poration (66). This confirms that we can use coarse-grained simulations to identify the lipid domains, which are the most likely to be porated. By backmapping (82) these regions to atomistic representation, we should be able to obtain a more accurate estimation of the kinetic parameters of eq. (2).

## 4 Conclusions

Pores in the plasma membrane can be formed under diverse physicochemical conditions. They can be formed in various physiological processes by pore-forming proteins, and when the membrane is subject to external mechanical or electromagnetic forces (83, 84). In this study we investigated pores formed by electric fields. Electroporation simulations of coarse-grained membranes mimicking realistic lipid composition of plasma membranes showed that pores preferentially form in domains enriched with polyunsaturated lipids and that pores avoid domains enriched with gangliosides, cholesterol, and fully saturated lipids. The density of polyunsaturated lipids is the most important feature governing the preferential pore location, regardless of the overall membrane composition and electric field polarity, as corroborated by machine learning methods. However, the poration kinetics does depend significantly on membrane composition and electric field polarity, as demonstrated by Bayesian survival analysis. The poration rate is higher under hyperpolarizing compared to depolarizing electric field and correlates with the ability of a membrane to expand its area under electric field. We envision that by combining atomistic and coarse-grained molecular dynamics simulations, Bayesian survival analysis and machine learning models, we will be able to improve existing kinetic models describing electroporation on the whole-cell level. Although we have focused on pores induced by electric fields, the findings are likely to be applicable to other ways of membrane poration, as we found that the lipid organization is much more important for poration than the electrical features of the membrane (charge, dipole angle).

## 5 Methods

### 5.1 Molecular dynamics simulations

#### System preparation

The starting point for our systems were the topology files for the coarse grained membranes used in the study of Ingolfsson et al. (36), available at https://bbs.llnl.gov/neuronal-membrane-data.html. The membranes are parametrized with the Martini 2.2 force field (34). We took the frames extracted after 80 μs of the simulation (the files confout-80us.gro). The original membranes were about 70 nm × 70 nm large. We cut four 30 nm × 30 nm pieces from the original membranes. After cutting, we removed an appropriate number of Na or Cl ions, such that the final system had zero net charge. We replaced the non-polarizable water model with its polarizable version to have a more accurate system representation for electroporation studies (85). We also added more water to each system, such that the simulation box in z direction was ~19 nm after equilibration. This procedure was done using functions from VMD (86), Gromacs (87), and custom scripts. In the end we had 8 membrane systems, four with lipid composition corresponding to the average plasma membrane (APM) and four corresponding to the brain plasma membrane (BPM) composition.

#### Equilibration and production runs

After preparing the new systems and topology files, we made two steps of system minimization and four steps of system equilibration following the equilibration protocol for Martini membranes from charmm-gui.org (88). We then ran a short 500 ns simulation using the reaction-field electrostatics followed by 50 ns equilibration using Particle Mesh Ewald (PME) electrostatics (89). Some of the PC lipid’s heads were constrained in their z position to reduce bilayer undulations, as done in the original publication (36). Other MD parameters were equal to the default parameters for simulations with Martini membranes from charmm-gui.org in April 2020: leap-frog integration of the equations of motion using 20 fs time step; plain cutoff of van der Waals and Coulomb interactions at distance 1.1 nm; relative dielectric constant *ε*_r_ = 2.5; temperature coupling using velocity rescaling with a stochastic term (90) with time constant 1 ps and temperature of 310 K; semi-isotropic pressure coupling using Parrinello-Rahman barostat (91) with time constant 12 ps and reference pressure of 1 bar, with the compressibility of the system set to 3e-4 bar^-1^. The trajectory was saved every 0.1 ns. All simulations were carried out with Gromacs 2019.4 (87).

#### Electroporation simulations

After equilibration, and for each of the 8 membranes, we ran multiple replicas of electroporation simulations where the membranes were exposed to an electric field of +127.7 mV/nm (60 replicas) or −127.7 mV/nm (60 replicas). No positional restraints were imposed in these simulations. The electric field was chosen such that we observed the formation of at least one pore within ~15 ns. Having a short poration time was important, because we aimed to map the local membrane features before poration to the likelihood of a poration event, which means that we needed to avoid considerable lateral lipid diffusion.

#### Additional simulations under non-porating electric field

For each membrane we also ran five 10-ns-long simulations under lower electric field strengths of 0, ±42.5, ±63.8, ±85.1, and ±106.3 V/nm. No positional restraints were imposed in these simulations. The initial coordinates (initial gro file) were the same as for electroporation simulations. With the exception of a few simulations at ±106.3 V/nm, these electric field strengths were too low to induce poration within 10 ns.

### 5.2 Pore localization

To determine the poration time and pore location in the electroporation trajectories we used a custom semi-automatic procedure with Python and MDAnalysis (https://www.mdanalysis.org/) (92). The procedure consisted of three main steps, as described below. For analysis we generally considered only the first poration event. Additional pores could form after the first one and all pores eventually expanded until destroying the membrane, as is usual for simulations under constant electric field (93, 94). We focused on the first poration event, as the local electric field changes after poration, changing the transmembrane voltage. In up to ~20% simulations, two or more pores formed practically simultaneously. In such a case, we considered all of these pores for analysis.

***Step 1 - Coarse Search:*** Start from frame 12 (since no pore formed before time 1.2 ns), search every 6 frames (i.e. 12, 18, 24 …). In each frame, divide the membrane into 100 small pieces. In each small piece, look for water molecules around the centre of the lipid bilayer. If there are more than 12 water molecules, stop and write down the frame (t) and the region (i,j), otherwise, continue the loop. This step is used to speed up and improve the accuracy of the algorithm. The condition of 12 water molecules ensured that only fully formed pores were considered for the second step. ***Step 2 - Precise Search:*** Start from frame t-5, search every frame until t (i.e. t-5, t-4, t-3, …). In each frame look for water molecules within the centre of the lipid bilayer in the regions (i-2, j-2), (i-2, j-1), …, (i+2, j+1), (i+2, j+2). If there are more than 4 (APM) or 6 (BPM) water molecules, check the following two frames. If there are water molecules also in the following two frames, return the poration time and the centre of mass of these water molecules as the pore location. Otherwise, continue the loop. ***Step 3 - Manual Check and Correction:*** The procedure failed for BPM, when the pore formed later than ~7 ns after the onset of the electric field, mainly because of the increase in BPM curvature with time. In this case, the poration time and location was determined by visualisation of the trajectory in VMD.

All poration times and pore locations were manually verified by extracting the porated frames from all trajectories and translating the systems in (x,y) such that the pore location moved to a predefined position. These extracted frames were then visualised in VMD.

### 5.3 Analysis of membrane properties

To determine local membrane properties we used the MemSurfer tool (37), available at https://github.com/LLNL/MemSurfer. MemSurfer is a 3D membrane surface fitting tool, which fits a surface to the inner and outer membrane leaflet and uses Delaunay triangulations and surface parameterizations to compute membrane properties of interest (Suppl. Fig. S2). Each vertex is mapped to the position of a lipid headgroup. The triangulated surface is then further smoothed (Suppl. Fig. S2d).

MemSurfer can be used to measure local membrane thickness, area per lipid, and mean curvature, among others. We added functions that analyse the dipole angle of zwitterionic lipids, the lipid tail order and the headgroup charge. The dipole angle for each zwitterionic lipid was computed as the angle between the local membrane normal (determined by MemSurfer) and the vector connecting two beads corresponding to the phosphate group and the choline or amine group (PO4 and NC3 or NH3). To determine lipid tail order, we first computed the angles between the local membrane normal and the vectors connecting all adjacent beads in the lipid tails. Then we computed the average of the cosine of all angles and determined the average order parameter as

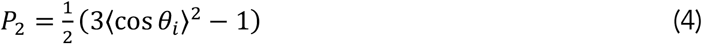

where the average goes over all bond angles in both lipid tails. This definition of the lipid order turned out to be more sensitive for separating values in porated and non-porated regions, compared to the more common definition 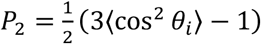 (36). We also extracted the type of lipid at a given vertex and the charge of this lipid. The scripts for running MemSurfer in this study, as well as the following analysis carried out in Matlab R2021a and described below, are available at https://github.com/learems/Electroporation-CGmem-MemSurfer.

MemSurfer returns the values of the above listed properties/features at vertices, which correspond to individual lipids’ headgroups. For a given trajectory frame, the values of most of the properties of interest like area per lipid, lipid order parameter, charge, …, can exhibit large variations among the adjacent vertices. Therefore, to extract the values at a selected (*x*_*p*_,*y*_*p*_) position, we used a Gaussian smoothing kernel, which determined a weighted average of the values *V*_*i*_ at all vertices within a radius 3*r*_*smooth*_ from the (*x*_*p*_,*y*_*p*_) position

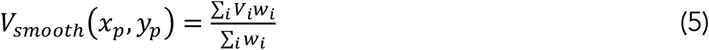

where the weights for a given adjacent vertex i are

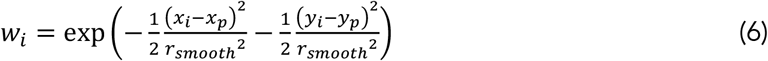

When determining the presence/density of lipid groups (PC, PE, SM, …), we assigned a value of 0 or 1 if a given lipid group was located at a given vertex. Gaussian smoothing was carried out in the same way as for other features. The same smoothing procedure was also done for plotting the surface plots in Fig. 5. We chose *r*_*smooth*_ = 1.5 nm, because this resulted in the best separation of the feature values in non-porated and porated regions.

The porated locations were defined as the locations at which a pore formed in any of the 60 electroporation simulations. For non-porated locations, we first divided the membrane into a grid with 31 by 31 points. We excluded all points which were within ~2 nm (6.7% of the box size) of any porated location. We also randomly excluded excess points such that the final number of non-porated locations was 300 (Fig. 3A).

The distances between histograms of feature values in porated and non-porated locations (Fig. 3B) were quantified by symmetrized version of the Kullback–Leibler divergence (38) after performing a kernel smoothing probability density estimate:

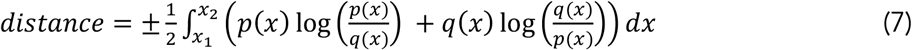

where *p*(*x*) and *q*(*x*) are the distributions of the data in porated and non-porated locations, respectively. The limits *x*_1_ and *x*_2_ were defined as the lowest and highest value of *x*, where either of the distributions fell under 1% of their peak value. In Fig. 3B the distances correspond to probability density estimates, which were obtained after averaging the value of a given feature and at a given point over both membrane leaflets. Distances computed for each leaflet separately are shown in Suppl. Fig. S9.

### 5.4 Machine learning methods

The values of all the features extracted in non-porated and porated locations from 101 frames of a 10-ns-long trajectory before electroporation were used to train three machine learning models with Python and scikit-learn library (95): random forest, support vector machine and multi-layer perceptron neural network. The input data contained the (x,y) locations on the membrane and the extracted membrane properties at those locations (i.e., features, denoted as X). The input data was separated into two classes with label Y = 1 for porated locations and Y = 0 for non-porated locations. As the number of non-porated locations (300) and porated locations (~60) were imbalanced, we used SMOTE in Python to randomly oversample the data in porated locations and balance them with the data in non-porated locations. After oversampling the data, we trained the above-listed machine learning models and used them to predict the pore locations. The accuracy of the prediction was evaluated as:

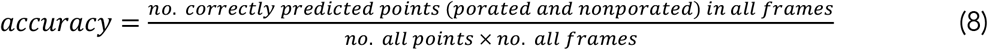

The codes used for this analysis are available at https://github.com/learems/Electroporation-CGmem-MachineLearning.

### 5.5 Bayesian survival analysis

To model the poration rates, we used Bayesian survival analysis. Given the non-exponential shape of the first poration times and the heterogeneity of poration rates between systems, it was crucial to use a model that allowed for an arbitrary time dependence of the rate and that was able to account for the system dependence. Cox’s proportional hazards model has these features and defines the event rate as

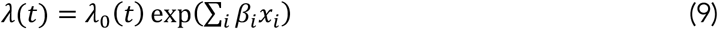

*λ*_0_(*t*) is constructed as a piecewise constant function defined in time intervals with endpoints: 0 ≤*s*_0_ ≤ … ≤ *s*_*N*_ such that *λ* _*0*_ (*t*) = *λ*_*j*_ for *s*_*j*_ ≤ *t* < *s*_*j*+1_. *x*_*i*_ are binary categorical variables such that if the poration time describes a system k, then *x*_*i*=*k*_ = 1 and *x*_*i*≠*k*_ = 0, i.e., one-hot-encoding representations of the system category. A good signal to noise ratio was found with 1.5 ns time intervals. *λ*_*j*_ were given independent priors in the form of gamma distributions with a shape parameter of 50 and a scale parameter of 10 ns^-1^. *β*_*i*_ were given normally distributed priors centred at the origin and with a standard deviation of 100. Both priors are fairly uninformative.

The Bayesian inference of the model parameters {*λ*_i_, *β*_*i*_} was done using as input features the first poration times and the system category (APM-dep, APM-hyp, BPM-dep, BPM-hyp) encoded in *x*_*i*_. In this way, the model was inferred on the electroporation events of all systems and *a posteriori* the model for a particular system i, *λ*_0_(*t*) exp(*β*_*j*_) was computed. Posterior predictive checks validated the quality of our models: the model can generate data that reproduces accurately the observed data (Suppl. Fig. S3).

To test the universality of *λ*_0_(*t*) and check the time-independence of *β*_*i*_, we inferred another model allowing *β*_*i*_ to be time dependent:

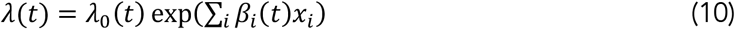

In this alternative model, *β*_*i*_(*t*) is also a piecewise constant function analogously to *λ*_0_(*t*). The heights of the steps of *β*_*i,j*_ for *s*_*j*_ ≤ *t* < *s*_*j*+1_ were given gaussian random walk priors: *β*_*i,j*_ is given a gaussian prior with standard deviation of 1 but centred on their previous *β*_*i,j*-1_.

Since we are using Bayesian statistics, the parameters of the model {*λ*_*j*_, *β*_*i*_} are treated as random variables whose distributions are inferred as a result. We can calculate their 94% credible intervals, i.e., the interval containing 94% of the probability density of the variable. Therefore, uncertainties are presented as the median of the variable ± the distance to the credible interval limits.

The models were inferred using PYMC3 (96) and follow a similar methodology to one of the library’s case studies (https://docs.pymc.io/notebooks/survival_analysis). The full details of the implementation are available at https://github.com/sperezconesa/electroporation_modeling.

## 6 Acknowledgements

This work was supported by grants from the Science for Life Laboratory, from the Gustafsson foundation, from Swedish Research Council (VR 2018-04905) to L.D., and by a synergy postdoc grant to L.D. and I.T. from KTH Royal Institute of Technology. The work was also supported by funding from the European Union’s Horizon 2020 research and innovation programme under the Marie Skłodowska-Curie grant agreement No. 893077 to L.R., and by funding from Slovenian Research Agency (ARRS) under project no. J2-2503. The simulations were performed on resources provided by the Swedish National Infrastructure for Computing at parallelldatorcentrum (PDC) Centre for High Performance Computing and at High Performance Computing Center North (HPC2N).

