## Supplementary Material for "Identification of electroporation sites in the complex lipid organization of the plasma membrane"

for

[1] KTH Royal Institute of Technology, Dept. Applied Physics, Science for Life Laboratory, Box 1031, 17121 Solna, Sweden
[2] University of Ljubljana, Faculty of Electrical Engineering, Tržaška 25, 1000 Ljubljana, Slovenia
[3] University of Chinese Academy of Sciences, Beijing 100049, China
[4] Authors contributed equally
* Corresponding authors

### S1. Membrane composition

The lipids are grouped into subtypes, depending on their headgroup type and tail saturation. Lipids are considered to be fully saturated, if they contain no double bonds in any of the tails. Lipids are considered to be monounsaturated, if they contain exactly one double bond in one or both of the tails. Lipids are considered to be polyunsaturated, if they contain at least two double bonds in one or both of the tails. Individual groups contain Martini lipids, which are listed in Table S1. The percentage of individual lipid groups in each of the four APM and BPM membranes is shown in Table S2. These percentages are also graphically presented in Fig. S1.

*Table S1: Separation of lipids into groups based on their headgroup type or tail saturation*

| *Group name* | *Abbrev.* | *Martini Lipids* |
|---|---|---|
| phosphatidylcholines | PC | DAPC, DOPC, DPPC, OIPC, OUPC, PAPC, PEPC, PFPC, PIPC, POPC, PUPC |
| phosphatidylethanolamines | PE | DAPE, DOPE, DUPE, OAPE, OIPE, OUPE, PAPE, PIPE, POPE, PQPE, PUPE |
| sphingomyelins | SM | BNSM, DBSM, DPSM, DXSM, PBSM, PGSM, PNSM, POSM, XNSM |
| gangliosides | GM | DBG1, DPG1, DXG1, PNG1, POG1, XNG1, DBG3, DPG3, DXG3, PNG3, POG3, XNG3, DBGS, DPGS, PNGS, POGS |
| ceramides | CE | DBCE, DPCE, DXCE, PNCE, POCE, XNCE |
| lysolipids | LPC | APC, IPC, OPC, PPC, UPC, IPE, PPE |
| diglycerides | DAG | PODG, PIDG, PADG, PUDG |
| phosphatidylserines | PS | DAPS, DOPS, DPPS, DUPS, OUPS, PAPS, PIPS, POPS, PQPS, PUPS |
| phosphatidylinositols | PI | POPI, PIPI, PAPI, PUPI |
| phosphatic acids | PA | POPA, PIPA, PAPA, PUPA |
| phosphatidylinositol phosphates | PIP | PAP1, PAP2, PAP3, POP1, POP2, POP3 |
| cholesterol | CHOL | CHOL |
| fully saturated tails | FS | DPPC, DBSM, DPSM, DXSM, PBSM, DPPS, DBCE, DPCE, DXCE, PPC, PPE, DBG1, DPG1, DXG1, DBG3, DPG3, DXG3, DBGS, DPGS |
| monounsaturated tails | MU | DOPC, POPC, DOPE, POPE, BNSM, PGSM, PNSM, POSM, XNSM, DOPS, POPS, POPI, POP1, POP2, POP3, POPA, PODG, PNCE, POCE, XNCE, OPC, PNG1, POG1, XNG1, PNG3, POG3, XNG3, PNGS, POGS |
| polyunsaturated tails | PU | OIPC, OUPC, PAPC, PEPC, PFPC, PIPC, PUPC, OAPE, OIPE, OUPE, PAPE, PIPE, PQPE, PUPE, OUPS, PAPS, PIPS, PQPS, PUPS, PAPI, PIPI, PUPI, PAP1, PAP2, PAP3, PAPA, PIPA, PUPA, PADG, PIDG, PUDG, APC, IPC, UPC, IPE, DAPC, DUPE, DAPE, DAPS, DUPS |

*Table S2: The total number of lipids and the percentage of individual lipid groups in each of the 8 membranes. The numbers correspond to inner / outer leaflet.*

| | *Average Plasma Membrane* | | | | *Brain Plasma Membrane* | | | |
|---|---|---|---|---|---|---|---|---|
| | *mem #1* | *mem #2* | *mem #3* | *mem #4* | *mem #1* | *mem #2* | *mem #3* | *mem #4* |
| *Total number of lipids in each leaflet* | | | | | | | | |
| *in / out* | 1503 / 1638 | 1564 / 1663 | 1571 / 1650 | 1553 / 1663 | 1728 / 1891 | 1740 / 1847 | 1770 / 1886 | 1764 / 1206 |
| *Percentage of lipid groups* | | | | | | | | |
| *CHOL* | 27.7 / 31.7 | 30.4 / 32.1 | 30.8 / 32.1 | 29.6 / 33.0 | 44.5 / 45.7 | 45.1 / 46.1 | 45.4 / 46.3 | 44.4 / 46.4 |
| *PC* | 17.9 / 36.4 | 18.9 / 34.3 | 15.9 / 34.7 | 17.8 / 34.3 | 12.4 / 23.0 | 12.4 / 23.6 | 14.8 / 25.8 | 14.2 / 21.9 |
| *PE* | 24.5 / 6.6 | 21.9 / 4.6 | 24.8 / 5.3 | 25.2 / 6.2 | 22.8 / 9.7 | 21.8 / 11.7 | 19.6 / 9.2 | 20.9 / 10.5 |
| *SM* | 10.6 / 18.7 | 8.9 / 19.5 | 9.7 / 19.6 | 8.6 / 19.1 | 2.1 / 8.9 | 2.1 / 9.0 | 2.3 / 9.2 | 2.8 / 8.5 |
| *GM* | 0.0 / 3.7 | 0.0 / 7.0 | 0.0 / 5.2 | 0.0 / 4.7 | 0.0 / 11.3 | 0.0 / 8.6 | 0.0 / 8.0 | 0.0 / 11.5 |
| *PS* | 11.3 / 0.0 | 10.7 / 0.0 | 9.6 / 0.0 | 10.6 / 0.0 | 9.8 / 0.0 | 11.0 / 0.0 | 9.1 / 0.0 | 8.7 / 0.0 |
| *PI* | 3.7 / 0.0 | 4.5 / 0.0 | 5.0 / 0.0 | 5.0 / 0.0 | 5.4 / 0.0 | 4.9 / 0.0 | 5.0 / 0.0 | 5.1 / 0.0 |
| *PIP* | 2.0 / 0.0 | 1.7 / 0.0 | 1.1 / 0.0 | 1.5 / 0.0 | 1.4 / 0.0 | 0.9 / 0.0 | 1.3 / 0.0 | 1.7 / 0.0 |
| *PA* | 1.6 / 0.0 | 2.1 / 0.0 | 2.0 / 0.0 | 0.9 / 0.0 | 0.3 / 0.0 | 0.6 / 0.0 | 0.5 / 0.0 | 0.2 / 0.0 |
| *LPC* | 0.0 / 1.5 | 0.0 / 0.8 | 0.0 / 1.4 | 0.0 / 1.0 | 0.5 / 0.3 | 0.2 / 0.5 | 0.5 / 0.6 | 0.3 / 0.3 |
| *DAG* | 0.5 / 1.0 | 0.3 / 0.5 | 0.4 / 0.8 | 0.3 / 1.0 | 0.4 / 0.2 | 0.3 / 0.1 | 0.8 / 0.3 | 1.0 / 0.1 |
| *CE* | 0.3 / 0.4 | 0.6 / 1.1 | 0.6 / 0.9 | 0.6 / 0.7 | 0.2 / 0.7 | 0.6 / 0.3 | 0.7 / 0.5 | 0.7 / 0.7 |
| *FS* | 6.6 / 12.2 | 4.4 / 14.8 | 5.3 / 14.7 | 4.8 / 14.0 | 5.1 / 22.4 | 5.2 / 19.3 | 6.1 / 19.8 | 5.8 / 20.6 |
| *MU* | 24.6 / 26.7 | 25.4 / 29.5 | 24.4 / 27.1 | 25.4 / 26.6 | 13.2 / 15.5 | 12.9 / 15.9 | 14.1 / 16.6 | 15.2 / 17.2 |
| *PU* | 41.1 / 29.4 | 39.8 / 23.6 | 39.5 / 26.2 | 40.2 / 26.5 | 37.2 / 16.3 | 36.8 / 18.7 | 34.4 / 17.2 | 34.5 / 15.8 |

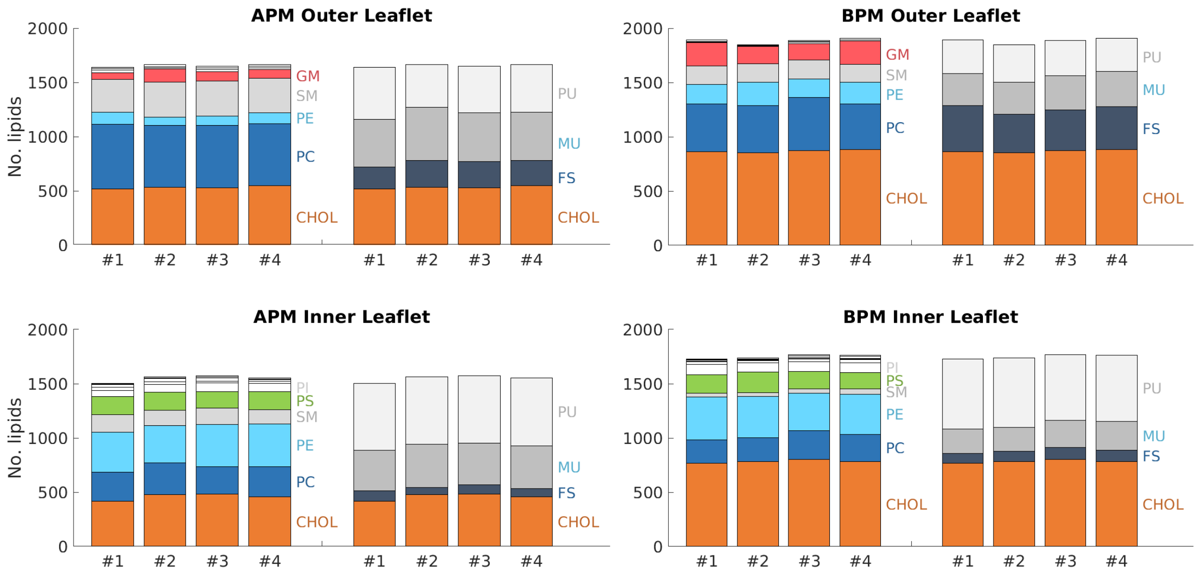

*Fig. S1:* Composition of the inner and outer leaflet of each of the four APM and BPM membranes.

### S2. Analysis of membrane properties

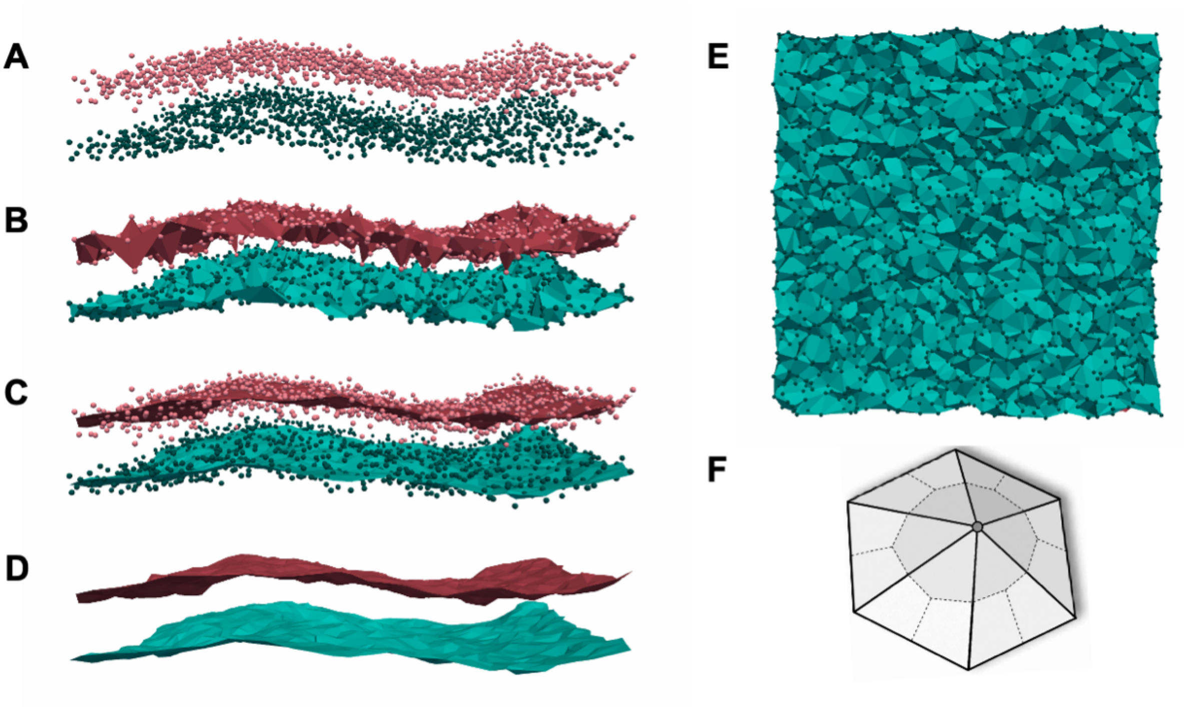

*Fig. S2: Membrane surface fitting with MemSurfer*. A) Points corresponding the lipid headgroups. B) Delaunay triangulations. C, D) Surface smoothed through a Poisson reconstruction approach. E) Top view of the triangulated surface. F) Determination of the area per lipid (shaded part).

### S3. Bayesian survival analysis

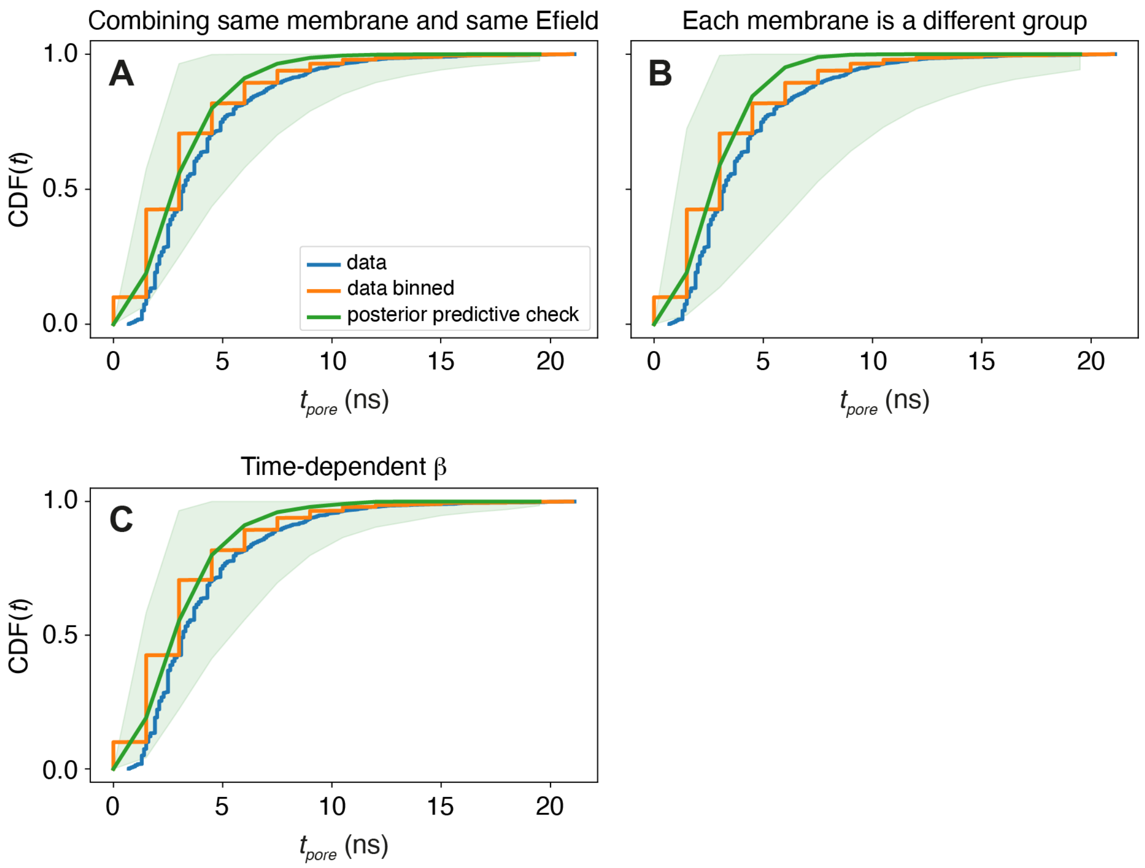

*Fig. S3:* Posterior predictive checks of the Bayesian inference procedures. We compare the observed data (blue), the observed data with the time discretized in order to be able to use stepwise defined functions in the model (orange), and the data that the model is able to generate (green). The agreement between the input data and the data generated by our modelling is satisfactory. The presented posterior predictive checks are done for A) combining the different membrane patches of the same membrane type and polarity in the same group; B) having each membrane fragment as a different group; and C) for the time dependent $\beta$ model.

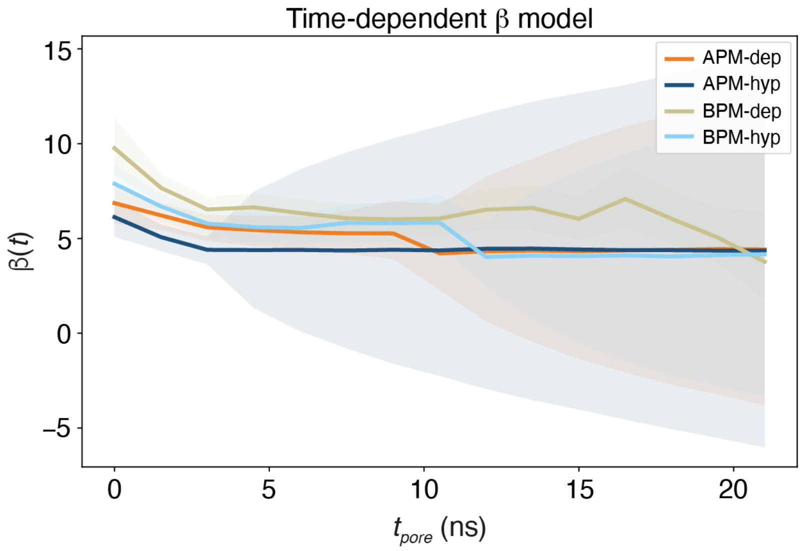

*Fig. S4:* Time evolution of the exp($\beta(t)$) for the time-dependent model.

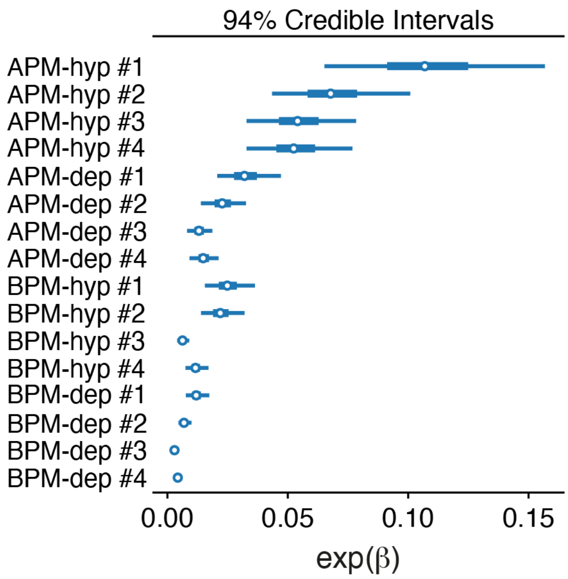

***Fig. S5:*** Credible intervals of exp($\beta_i$) separately for each of the 8 membranes considered and both electric field polarities.

### S4. Membrane response to nonporating electric fields

Upon exposure to electric field, the membranes increase their area. The relative increase in the membrane area in steady state is proportional to $E^2$ (Fig. S6), as expected for Maxwell stress. The slopes of the of the least-square fitted lines in Suppl. Fig. S6 were used to plot Fig. 5E.

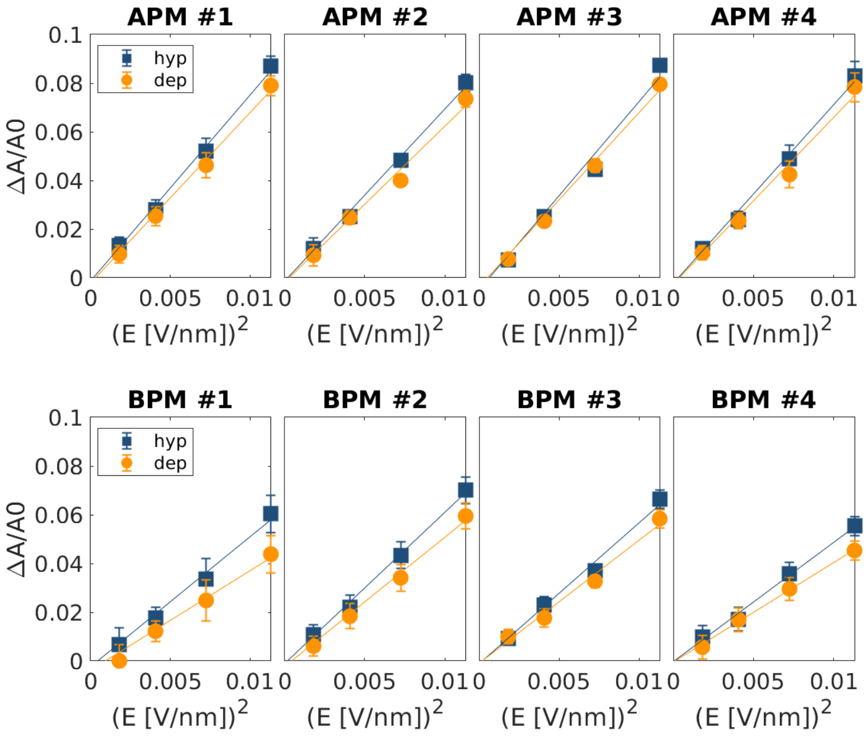

***Fig. S6: Relative change in the membrane area under nonporating electric fields.*** The relationship between the steady state relative change in the membrane area $\Delta A/A_0$ and $E^2$. Each graph corresponds to one of 8 simulated membranes; hyp=hyperpolarization, dep=depolarization.

### S5. Histograms

APM - depolarization - inner leaflet

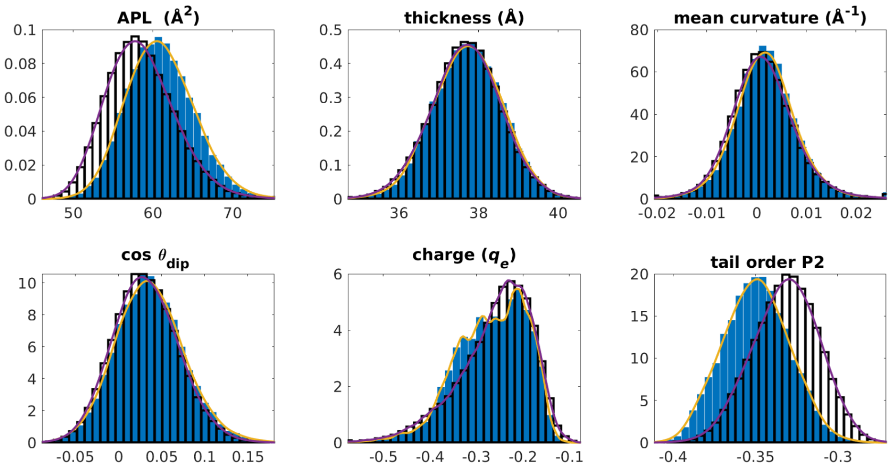

APM - depolarization - outer leaflet

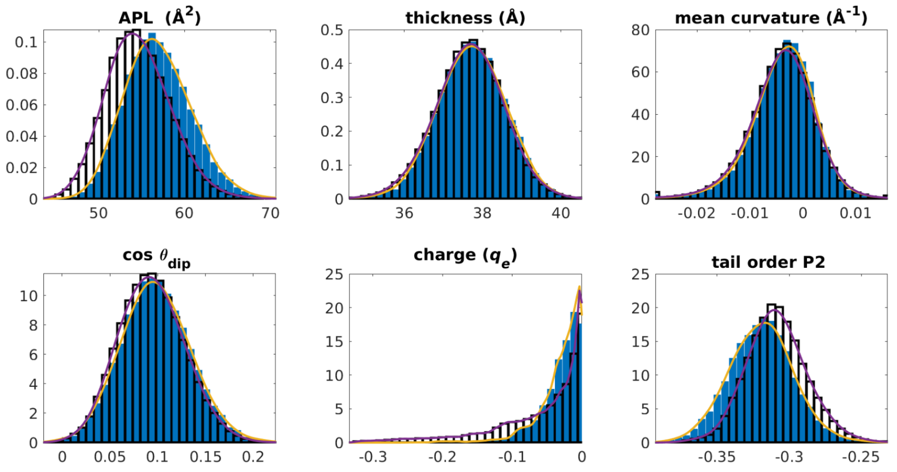

APM - hyperpolarization - inner leaflet

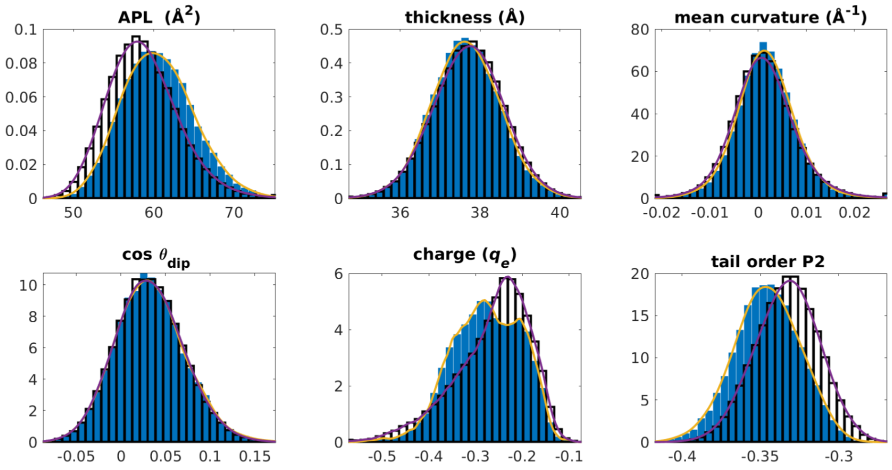

APM - hyperpolarization - outer leaflet

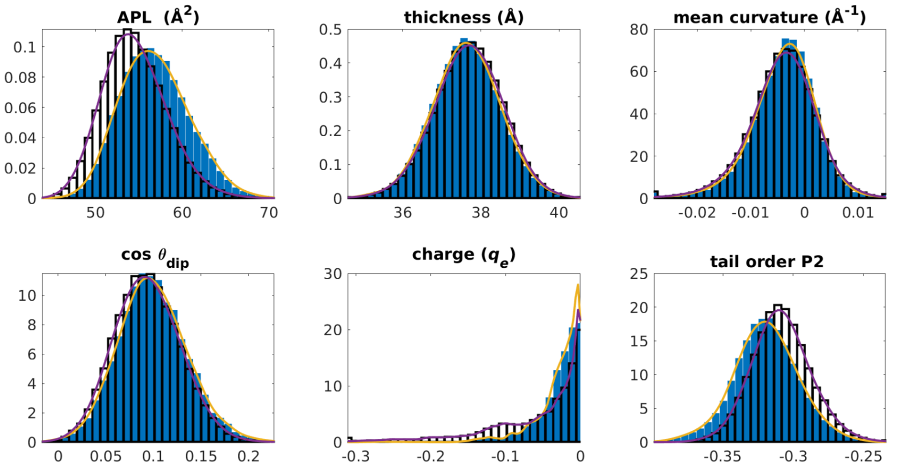

*Fig. S7: Histograms for APM.* Histograms showing the values of tested features in nonporated (empty bars) and porated (blue bars) locations. The purple and yellow solid lines show kernel smoothing density estimates. The title above each group of graphs indicates the dataset for which the histograms are shown.

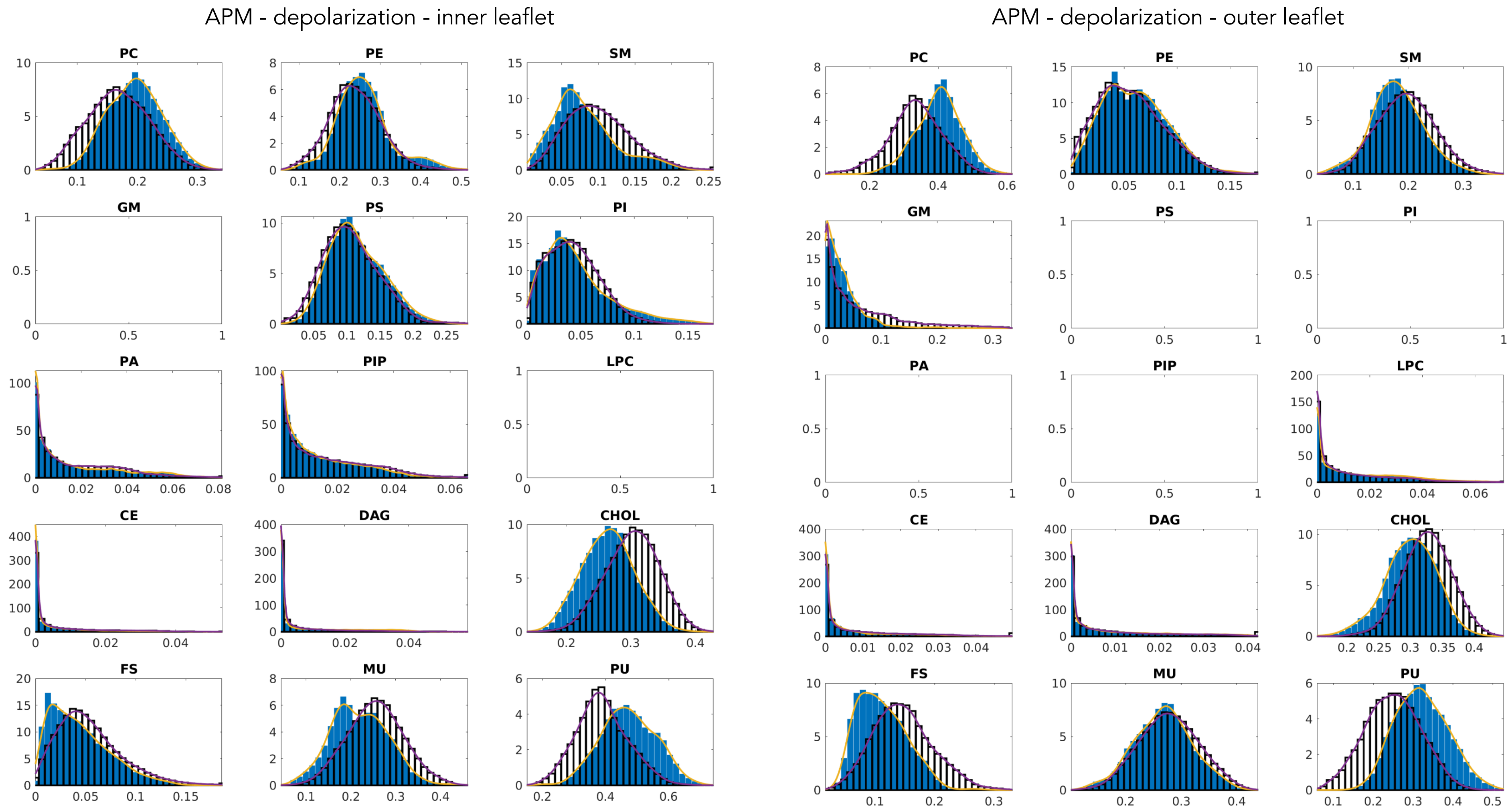

*Fig. S7: Histograms for APM - continued.*

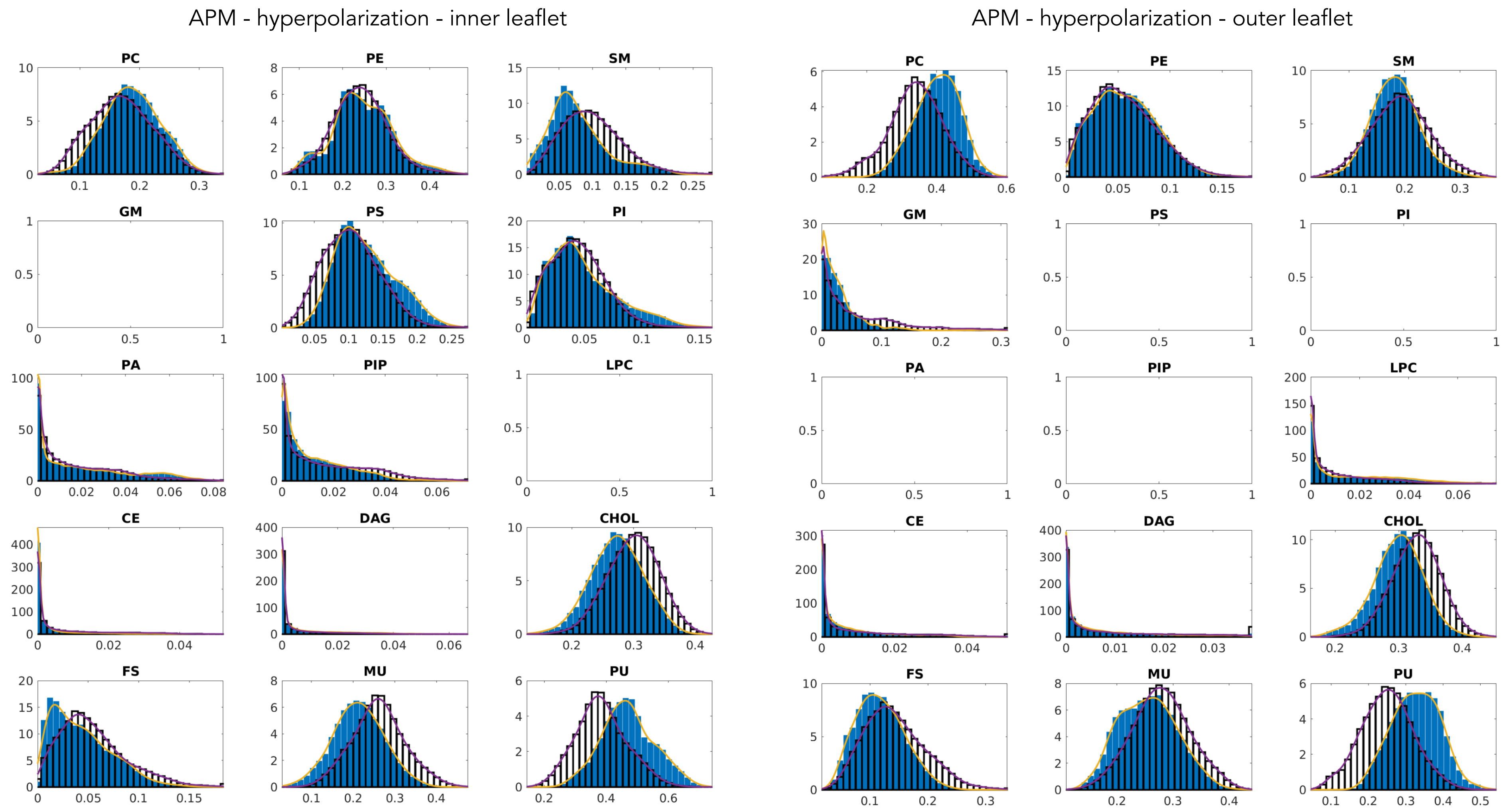

*Fig. S7: Histograms for APM - continued.*

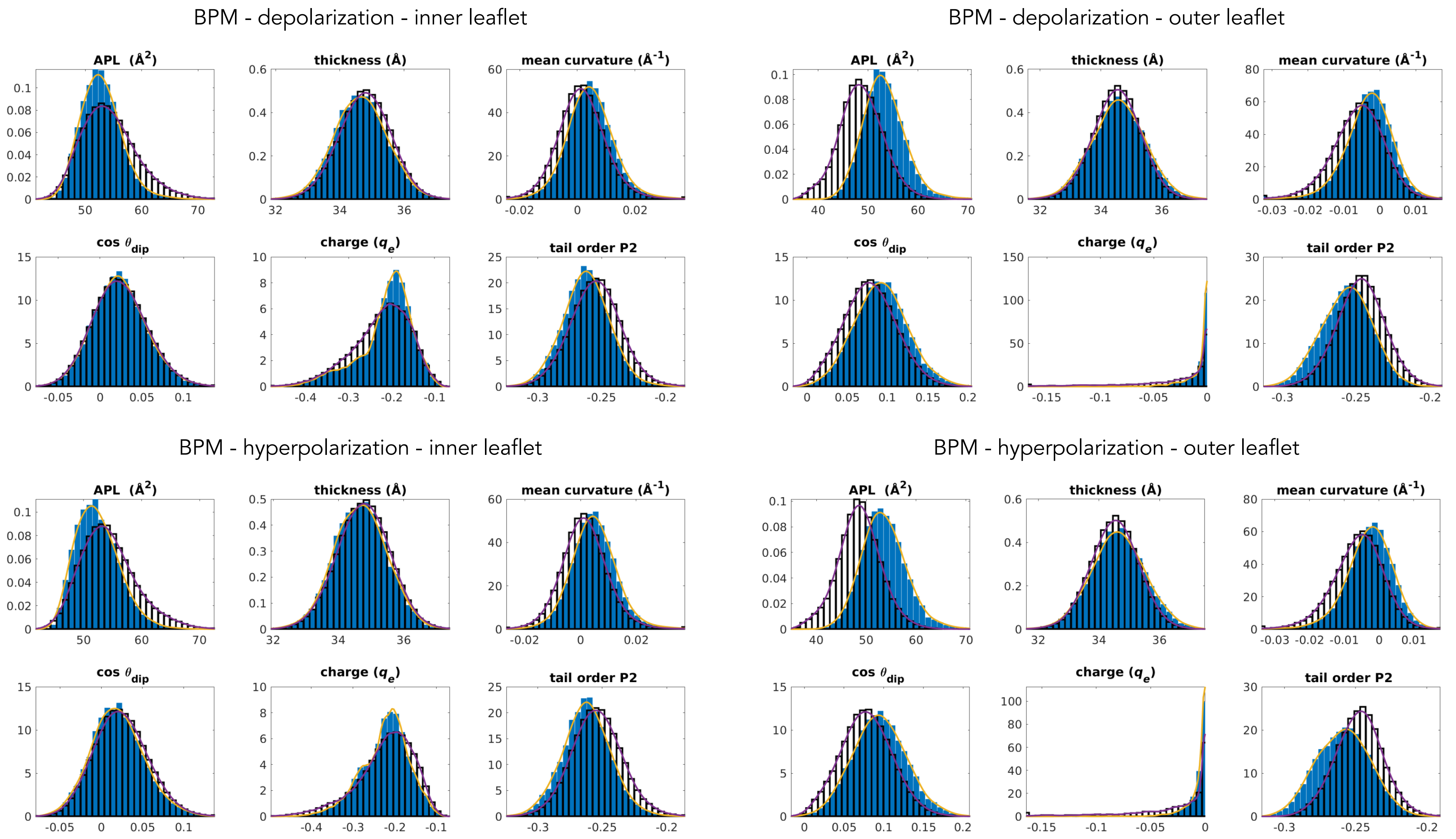

*Fig. S8: Histograms for BPM.* Histograms showing the values of tested features in nonporated (empty bars) and porated (blue bars) locations. The purple and yellow solid lines show kernel smoothing density estimates. The title above each group of graphs indicates the dataset for which the histograms are shown.

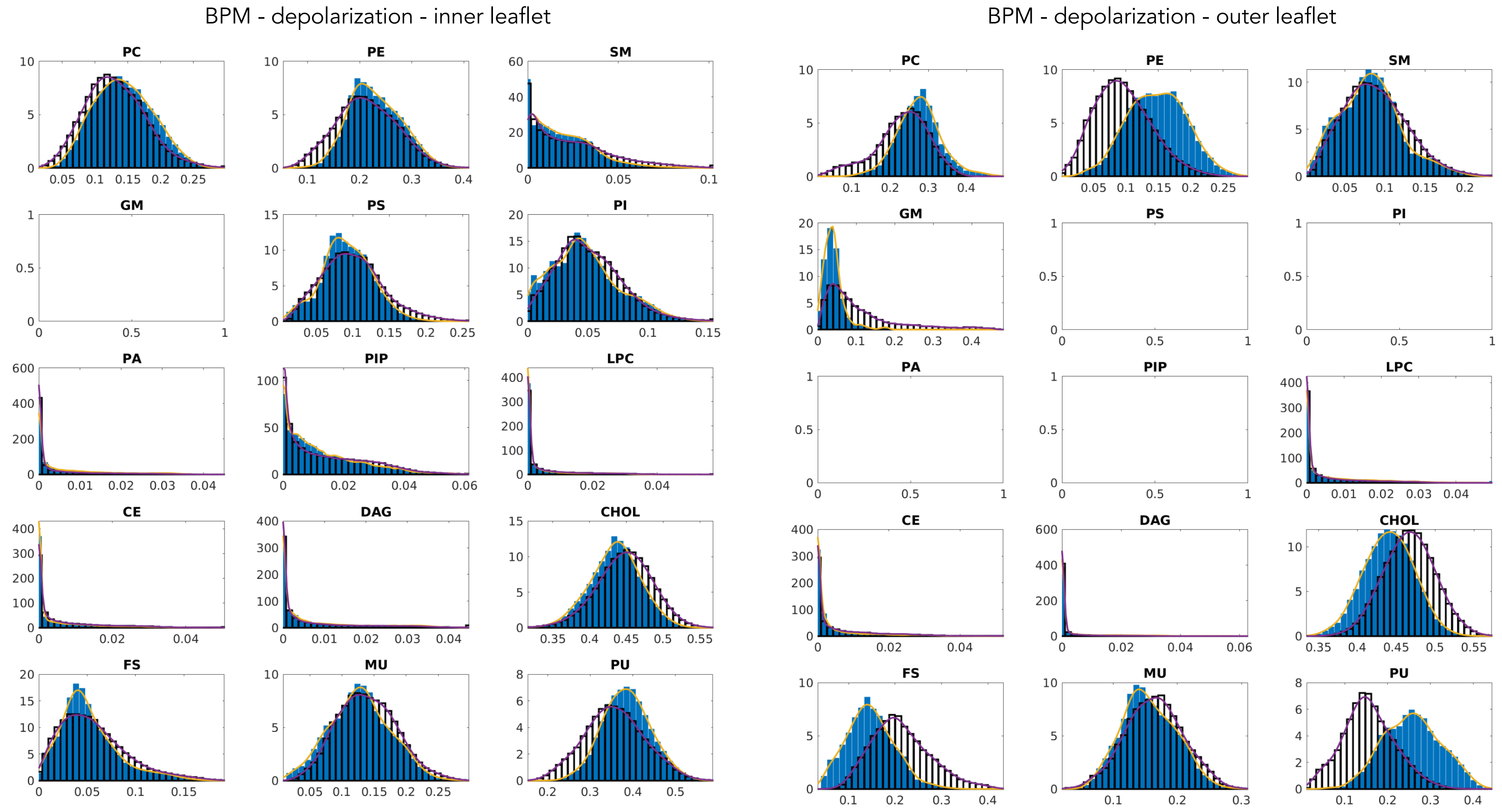

*Fig. S8: Histograms for BPM - continued.*

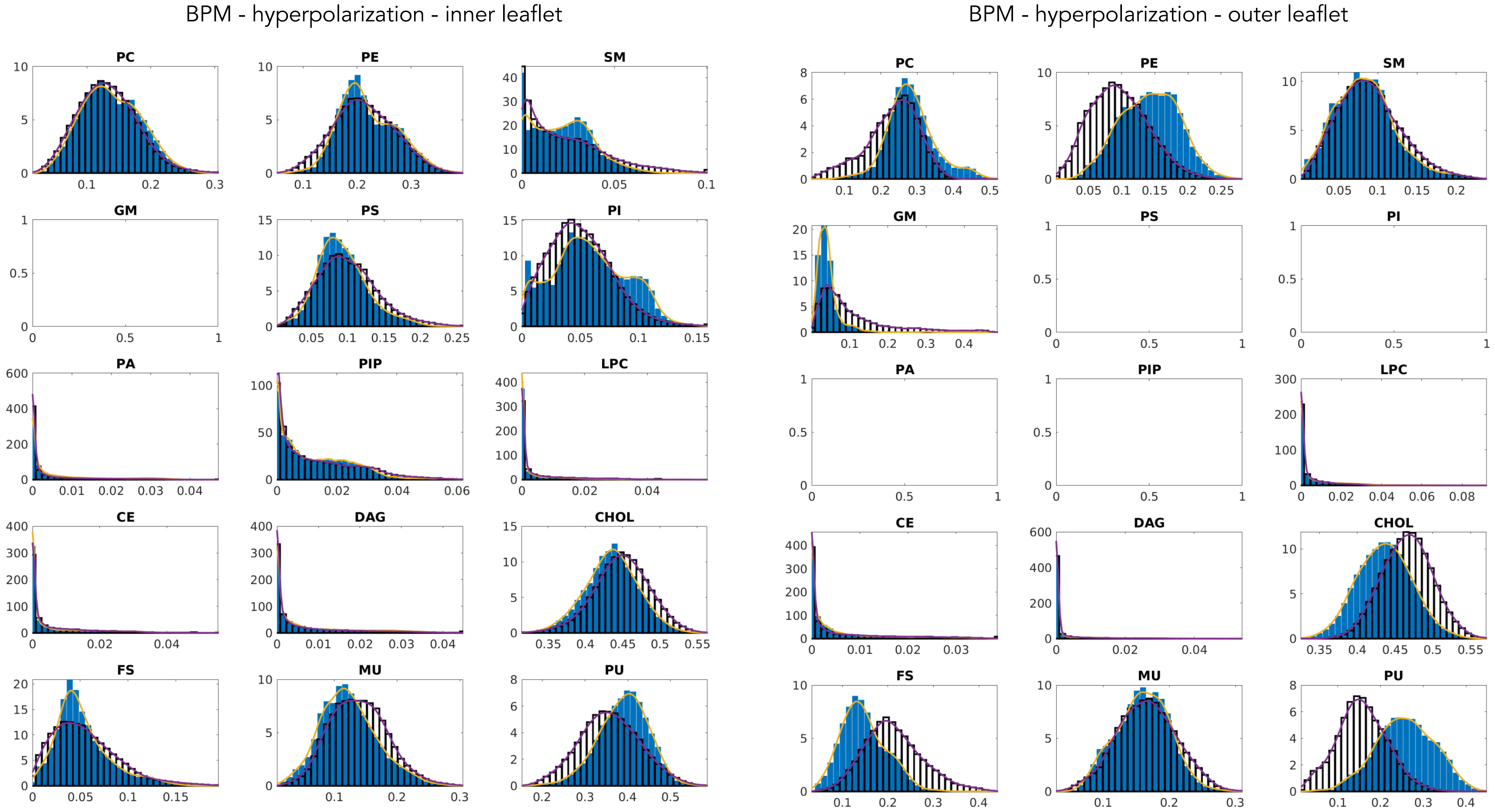

*Fig. S8: Histograms for BPM - continued.*

### S6. Distances between probability density estimates (KL divergence)

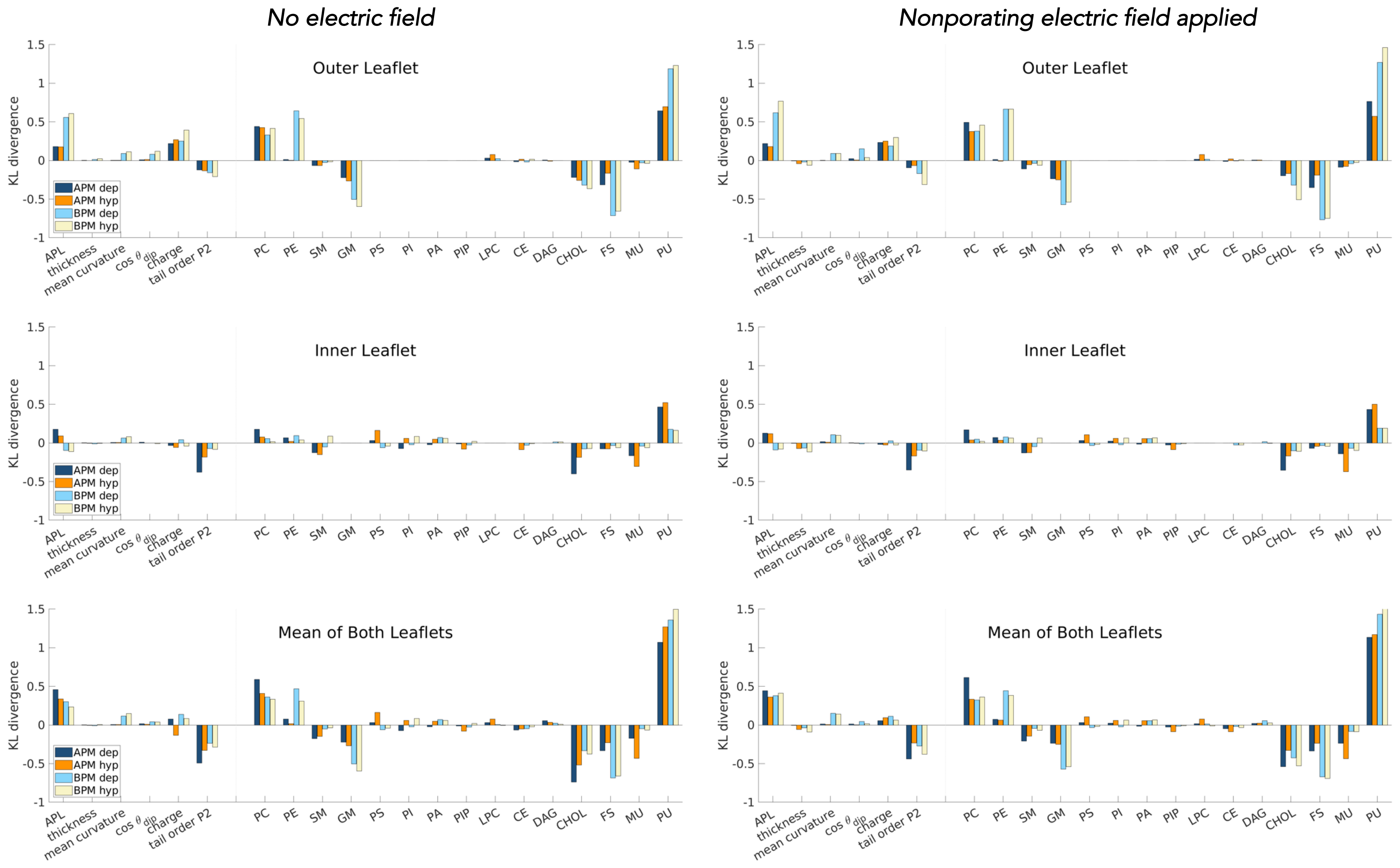

***Fig. S9: Distances between histograms.*** Distances between probability density estimates of individual features in porated and nonporated locations, calculated according to eq. (2). Graphs are shown separately for the outer and inner leaflet and for the mean of both leaflets, see the titles above each graph. The bigger the bar, the more the feature influences poration. Positive and negative bars shows whether a feature favours or disfavours poration, respectively. The graphs on the left side correspond to values extracted from a 10-ns-long trajectory before electric field application. The graphs on the right side correspond to values extracted from a 10-ns-long trajectory, where a non-porating electric field of +106.3 mV/nm (analysis for depolarization) or –106.3 mV/nm (analysis for hyperpolarization).
